# Identification and functional characterization of two novel mutations in *KCNJ10* and *PI4KB* in SeSAME syndrome without electrolyte imbalance

**DOI:** 10.1101/506949

**Authors:** Ravi K Nadella, Anirudh Chellappa, Anand G Subramaniam, Ravi Prabhakar More, Srividya Shetty, Suriya Prakash, Nikhil Ratna, VP Vandana, Meera Purushottam, Jitender Saini, Biju Viswanath, PS Bindu, Madhu Nagappa, Bhupesh Mehta, Sanjeev Jain, Ramakrishnan Kannan

**Affiliations:** Department of Psychiatry at National Institute of Mental Health and Neurosciences, Bangalore, India; Department of Neurology at National Institute of Mental Health and Neurosciences, Bangalore, India; Department of Biophysics at National Institute of Mental Health and Neurosciences, Bangalore, India; Department of Neuroimaging and Interventional radiology at National Institute of Mental Health and Neurosciences, Bangalore, India; Department of Speech Pathology and Audiology at National Institute of Mental Health and Neurosciences, Bangalore, India; National Centre for Biological Sciences, Tata Institute for Fundamental Research, Bangalore, India

**Author notes:** Address for correspondence Dr. Ramakrishnan Kannan, PhD, National Institute of Mental Health and Neurosciences (NIMHANS), Bengaluru, Karnataka, India PIN: 560029.

## Abstract

Dysfunction in inwardly-rectifying potassium channel Kir4.1 has been implicated in SeSAME syndrome, an autosomal-recessive (AR), rare, multi-systemic disorder. However, not all neurological, intellectual disability and comorbid phenotypes in SeSAME syndrome can be mechanistically linked solely to Kir4.1 dysfunction. We therefore performed whole exome sequencing and identified additional genetic risk-elements that might exert causative effects either alone or in concert with Kir4.1 in a family diagnosed with SeSAME syndrome. Two variant prioritization pipelines based on AR inheritance and runs of homozygosity (ROH), identified two novel homozygous variants in *KCNJ10* and *PI4KB* and five rare homozygous variants in *PVRL4, RORC, FLG2, FCRL1, NIT1* and one common homozygous variant in *HSPA6* segregating in all four patients. The novel mutation in *KCNJ10* resides in the cytoplasmic domain of Kir4.1, a seat of phosphatidyl inositol bisphosphate (PIP2) binding. The mutation altered the subcellular localization and stability of Kir4.1 in patient-specific lymphoblastoid cells (LCLs) compared to parental controls. Barium-sensitive endogenous K^+^ currents in patient-specific LCLs using whole-cell patch clamp electrophysiology revealed membrane depolarization and defects in inward K^+^ ion conductance across the membrane, thereby suggesting a loss-of-function effect of *KCNJ10* variant. Altogether our findings implicate the role of new genes in SeSAME syndrome without electrolyte imbalance and thereby speculate the regulation of Kir4.1 channel activity by PIP2 and integrin-mediated adhesion signaling mechanisms.

## Introduction

Channelopathies are a heterogeneous group of disorders resulting in dysfunction of ion channels. They disrupt the brain function resulting in seizures and developmental delay [1, 2, 3, 4, 5, 6, 7, 8]. The cells of central and peripheral nervous system contain a plethora of ion channel proteins which interact with multiple signaling pathways linking channel physiology to neuronal differentiation, axonal integrity and cell migration [6, 7, 9, 10]. Nevertheless, not all phenotypes manifested in a syndromic disorder can be attributed to monogenic variants in membrane ion channels [11]. Therefore, for a complete molecular understanding of channelopathies, it is imperative to focus on other classes of risk-associated rare variants especially in minor genes which modifies the effect of major gene mutations. Such an approach for SeSAME syndrome, a rare autosomal recessive, multisystemic neuropsychiatric illness has not been addressed and will greatly benefit to understand the aetiology of Kir4.1 channel dysfunction that will ultimately inform treatment.

SeSAME syndrome (OMIM#612780), characterized by **S**eizures, **S**ensorineural deafness, **A**taxia, **M**ental retardation and **E**lectrolyte imbalance, otherwise known as EAST (**E**pilepsy, **A**taxia, **S**ensorineural deafness, **T**ubulopathy) syndrome is predominantly caused by homozygous or compound heterozygous mutations in *KCNJ10* gene [12, 13] encoding Kir4.1, an inwardly rectifying potassium channel. Till date, 21 mutations from 27 patients have been reported, of which 11 were from consanguineous unions [14]. Dysfunction of Kir4.1 has been associated with other neurodegenerative conditions like idiopathic epilepsy [15], autism spectrum disorder with seizures [16, 17], Huntington’s disease [18], multiple sclerosis [19] and Rett syndrome [20]. Several modern-day mammals like Jack Russell Terriers, Belgian Shepherd dogs [21] and Malinois dogs [22] experienced SeSAME-like phenotype with *KCNJ10* mutations.

Kir 4.1 channels display greater inward K^+^ flow at negative resting membrane potential to equilibrium potential for K^+^ (*E*k), while at more positive membrane potentials, outward flow of K^+^ is inhibited by intracellular Mg^2+^ and polyamines [23]. Depending on tissue localization and assembly of Kir4.1 subunit, these channels exhibit distinctive physiological properties [24]. Kir4.1 channel play conspicuous roles in a spectrum of biological contexts like maintenance of resting membrane potential [25], facilitation of glutamate uptake [26], potassium siphoning by glial cells [27, 28], cell volume and peak strength regulation of motor neurons [10], axonal integrity through myelination by oligodendrocytes [6, 7, 29] and cell migration [9]. How Kir4.1 drives specific downstream signaling during disease manifestation in SeSAME syndrome requires us to understand the plethora of modifiers. Moreover, the activation of Kir4.1 depend inherently on factors like cellular milieu, presence of auxiliary subunits and formation of subunits for heterooligomeric assembly in cell type of choice [27]. To address these issues and to identify other genetic associative elements with *KCNJ10*-mediated SeSAME pathogenesis, we performed whole exome sequencing and functional characterization of pathogenic *KCNJ10* variant in patient-specific lymphoblastoid cells which harbours the spectrum of risk variants.

Whole exome sequencing analysis of four patients and two unaffected parents identified a novel missense mutation in *KCNJ10*, a candidate gene in SeSAME syndrome. In addition, using two independent variant prioritization pipelines, we isolated variants in other minor genes which are known to be involved in pathways that regulate Kir4.1 signaling in different biological contexts. Along with *KCNJ10*, our pipeline also identified novel variants in the following genes; *PIK4B* (PIP2 signaling), *PVRL4* (cell adhesion signaling), *HSPA6* (ER-protein trafficking) and *NIT1* (apoptosis). Finally, we validated the impact of *KCNJ10* variant in inward-rectification of K^+^ current using patient-specific LCLs. The variant is localized in a stretch of conserved residues required for PIP2 binding which is juxtaposed at the junction of transmembrane and cytoplasmic domain. Functionally, the variant alters its protein localization, accumulates in the cytoplasm, depolarizes the membranes and inhibits inward-rectification of K^+^ currents in patient LCLs.

## Materials and Methods

### Patient recruitment, genomic DNA isolation and generation of lymphoblastoid cells

Blood samples collected from ten participants [unaffected parents, (n=4), and affected off springs, (n=6)]after receipt of informed consent were recruited at the National Institute of Mental Health and Neurosciences under aseptic conditions following guidelines established by Institutional Human Ethics Committee (IHEC) and Institutional Stem Cell committee (ISCC). The participants were referred for biochemical evaluation and selected for further analysis by presence of clinical features like seizures, ataxia, mental retardation, hearing impairment. Genomic DNA was isolated from blood samples of all participants using NucleoSpin^®^ Blood L (Macherey-Nagel GmbH & Co. KG) for whole exome sequencing (WES). Peripheral blood mononuclear cells (PBMNCs) was isolated from whole blood of ten individuals and transformed by Epstein Barr virus (EBV) using standard protocol [30] to generate lymphoblastoid cell lines (LCLs). The six LCLs suspensions were cultured in medium supplemented with RPMI-1640 (HiMedia AL060A), 20% fetal bovine serum (Thermo Fisher Scientific 16000-044), 1% penicillin/streptomycin (Thermo Fisher Scientific 15140-122) and maintained at 37°C with 5 % CO_2_ in a humidified atmosphere. The LCLs were further screened for karyotype abnormalities using G- banding approach and sample identity confirmation was done by STR profiling [GenePrint® 10 System (Promega)].

### Whole exome sequencing, variant calling, quality check and annotation

DNA library was prepared using Nextera Rapid Capture and Expanded Exome Kits. The library was further subjected to WES, performed on Illumina Hi-Sequencer to generate pair-end reads (150bp*2). We followed whole exome sequence analysis pipeline used by [31]. FastQC (v0.11.5) (http://www.bioinformatics.babraham.ac.uk/projects/fastqc) was used for the quality of raw reads, which examine per base and per sequence quality scores, per base and per sequence GC content, per base N content and sequence length distribution. Prinseq-lite-0.20.4 tool was used to trim poor quality region (http://prinseq.sourceforge.net/) and adapterremoval-2.1.7 was used to remove adapter contamination in raw reads. Filtered reads with a quality score (Q)>20 were aligned to the human reference genome hg19 (GRCh37) using BWA (v0.5.9). SAM to BAM conversion and sorting were done with Samtools 1.3 tool (https://sourceforge.net/projects/samtools/files/samtools/1.3/). Then the PCR duplicates were removed using PICARD tools (v1.96) (https://broadinstitute.github.io/picard/) and the INDELS were realigned using GATK (v3.6). The BAM alignment was subjected to QC using Qualimap (v2.2). VarScan (v2.3.9) (Coverage=8, MAF>=0.25, *p*-value<0.001) was used to call for SNPs and INDELS. The quality of VCF file was checked using RTG tools 3.7.1 (https://github.com/RealTimeGenomics/rtg-tools/releases). All samples annotation was performed using ANNOVAR tool. Population controls (n=7) representing three religiousgroups (Group A, B, and C) matched for age, sex and ethnicity, were obtained from INDEX-db [32]. All controls passed the age of risk i.e., 45 years, for neuropsychiatric illnesses, except for the outbred Parsi (religious group 3) individual (age=26), who was included as an outlier. All the controls were of southern Indian ethnic origin except for the Parsi. To validate *KCNJ10* variant identified by whole exome sequencing, we performed Sanger validation using the following gene specific primers: Forward (CATTCGTTTCAGCCAGCATGC) and Reverse (TCAGACATTGCTGATGCGCA).

### Assessing runs of homozygosity (ROH)

Exome-wide F-statistics was calculated using the --het option in *vcftools* (v0.1.5), for every sample to investigate whether levels of heterozygosity differed between the affected siblings, unaffected parents and population controls. Runs of homozygosity (ROH) was detected in all samples using --homozyg option in PLINK (v1.9) [33]. The minimum length for a tract to qualify as ROH was set to 500kb and the minimum number of variants constituting an ROH was set to 100. A maximum of 3 intervening heterozygous variants were allowed within a ROH window. ROH density was set to default i.e., an ROH must have at least one variant per 50kb, on an average. The centromeric, X, Y and mitochondrial variants were ignored during this analysis. The stretches that were shared between all the affected individuals but not observed in either of the parents or the population controls were thus notified as ROH_affected_, which were identified by using a combination of *intersect* and *subtract* functions in *bedtools* (v2.22). The variants were annotated using variant effect predictor (VEP GRCh37).

### Whole-cell patch clamp electrophysiology

For electrophysiology studies, LCLs from a healthy wild type control, six participants from SeSAME like family described in this study were used. The LCLs were dissociated to single cells and plated on glass cover slips coated with poly-D-lysine (Millipore, A003M EMD) and incubated for half an hour at 37°C with 5% CO_2_ in a humidified atmosphere before recordings. Whole cell patch clamp recordings were configured following which the membrane potential (Vm) of LCLs was measured. A pulse protocol was applied with Vm held at resting membrane potential and then stepped to test potentials between −120mV to 40mV in 10mV steps for 140ms. A single electrode was used to measure membrane current (nA) by whole cell patch clamp technique. Intracellular voltage-clamp recordings and positioning of perfusion micropipette were done using two Narashige hydraulic micromanipulators (MNW-203, Narashige Japan). Recording pipettes (tip resistance 4-6MΩ) were filled with intracellular solution containing 120mM potassium D-gluconate (G4500, Sigma), 1mM MgCl_2_, 15mM KCl, 1mM CaCl_2_, 10mM EGTA, 10mM HEPES (pH 7.2). After obtaining whole-cell mode, access resistance was 10-15 MΩ. The extracellular recording solution contained 130mM NaCl_2_, 3mM CaCl_2_, 2.5mM MgCl_2_, 15mM HEPES (pH 7.4). In experiments, where LCLs were perfused with high extracellular K^+^, concentration of KCl varied from 5-20 mM while that of NaCl was decreased to 110mM to adjust osmolarity. Recordings in LCLs were performed using an HEKA triple patch clamp amplifiers (EPC 10 USB) at room temperature (RT). To determine specificity of Kir4.1 current, 110μm/L BaCl_2_ was used and to block endogenous Cl^-^ currents, 150μm/L niflumic acid was used in the bath solution. The pClamp 9 (Axon Instruments) software package was used for data acquisition and analysis. For statistical analysis we used GraphPad Prism (San Diego, USA). To chose between parametric or non-parametric tests for normality criteria, Shapiro-Wilk estimator was used. For data sets with small N, non-parametric test was used to avoid possible type II errors. Mean differences were statistically evaluated using ANOVA with Levene’s homogeneity of variances test and pairwise comparisons were made using Turkey adjustment. Non-parametric *k* independent Kruskal-Wallis test was applied with Bonferroni correction to compare the differences among means. Error bars represent ±S.E.

### Immunofluorescence and western blotting

The LCLs were fixed using 4% paraformaldehyde (Sigma, PFA: P6148) in phosphate buffered saline (PBS) for 20 min at RT. Cells were permeabilized using 0.2 % Triton X-100 (Sigma, T8787) for 10 min and were washed twice with PBS. Following permeabilization cells were blocked for 1h using 2% bovine serum albumin (BSA) in PBST (PBS containing 0.05% tween 20; Sigma, P2287). Primary antibody against hKir 4.1 (1:100, Novus biologicals, NBP1-20149) was incubated overnight at 4°C in block solution. Cells were washed twice with PBST followed by 1h incubation at RT with anti-rabbit Alexa Fluor^TM^ 488 (1:200; Thermo Fisher Scientific, A11001) and Alexa Fluor^TM^ 568 phalloidin (1:200; Thermo Fisher Scientific, A12380). Following incubation cells were washed twice with PBST and incubated with DAPI (1:10000; Thermo Fisher Scientific, 62248) for 10 min at RT. The cells were washed twice with PBS and mounted using Vectashield antifade mounting medium (H-1000: Vector labs). Optical *z*-sectioning at 0.2 μM intervals was done using Plan-Apochromat 63x/1.40 oil objective in Zeiss Axio Observer 7 with Apotome 2 feature and Axiocam 702 monochrome camera (Carl Zeiss, Germany). Signal-to-noise ratio was improved using in-built Zeiss deconvolution module and MIP projections of 2-3 *Z*-stacks are presented here. Representative images reported here are from three independent experiments. For quantitative measurements, deconvoluted Z-stacks were first blinded before analysis. 3D surface rendering plugin in Imaris software is used to reduce signal-noise ratio to measure Kir4.1 punctate distribution between cytoplasm and nucleus. The respective numbers were normalized against cytoplasmic space marked by F-actin and nuclear space by DAPI signals.

LCLs suspension of all six participants were cleared by centrifugation (1500 rpm for 3 min) to remove culture media. RIPA lysis buffer containing phosphatase and protease inhibitor cocktails (EDTA-free, ab201120) was used to lyse the cells and total protein was isolated. Bradford assay was used to measure the concentration of the protein. All six samples (20 ug protein /lane) were resolved using 10% SDS-PAGE, transferred to PVDF membrane and probed with anti-Kir4.1 protein (NBP1-20149) and β-actin (A5441) as loading control. Target protein bands detection was done in Gel Documentation system (Syngene: chemiXX9) using Super signal West Pico Chemiluminescent substrate (Thermo Scientific, #34077) and densitometric quantitation assessed using Image Studio Lite v5.2 (LI-COR Biosciences).

## Results

### Clinical features of a family with SeSAMEsyndrome

Six affected patients,born through two consanguineous unions, were identified from the relatives of an index patient (IV.2) who developed tonic-clonic seizures, ataxiaand developmental delay (Fig. 1a). The clinical features were broadly similar to SeSAME syndrome but without electrolyte imbalance (Table 1). The cerebellar symptoms (gait ataxia, intentional tremors and dysdiadochokinesia) were manifested from early childhood. The gait ataxia was progressive in nature, resulting in severe disability and later being confined to wheel chairs [IV. 2-5]. Dysmorphic facies, dysarthria, brisk deep tendon reflexes (DTRs), bilateral ankle clonus and an extensor Babinski response were evident in all of them. All the patients showed certain characteristic dysmorphic facial features like prominent supraorbital ridges, thick eyebrows, deep set eyes, epicanthal fold, low set ears, prominent antihelix, prominent nasal tip and thick lips (Fig. 1b). Behavioural abnormalities like stereotypies, hyperactivity, anger outbursts and psychotic symptoms were also observed (Table 1). They also had hearing impairment, and audiometry measures revealed bilateral mild to severe sensory neural hearing loss. Motor nerve conduction velocities from patients (V.1-2) were normal. The EEG from patients (V.1-2) showed generalised seizure discharges before treatment (Fig. 1c), which became normal after treatment with anti-epileptic drugs. The other four members (IV.2-5) remained seizure free for several years on medication.MRI from IV.2 showed enlarged basal ganglia and cerebellar atrophy (Fig. 1d). The remaining members of the family were clinically unaffected.

**Figure 1.**
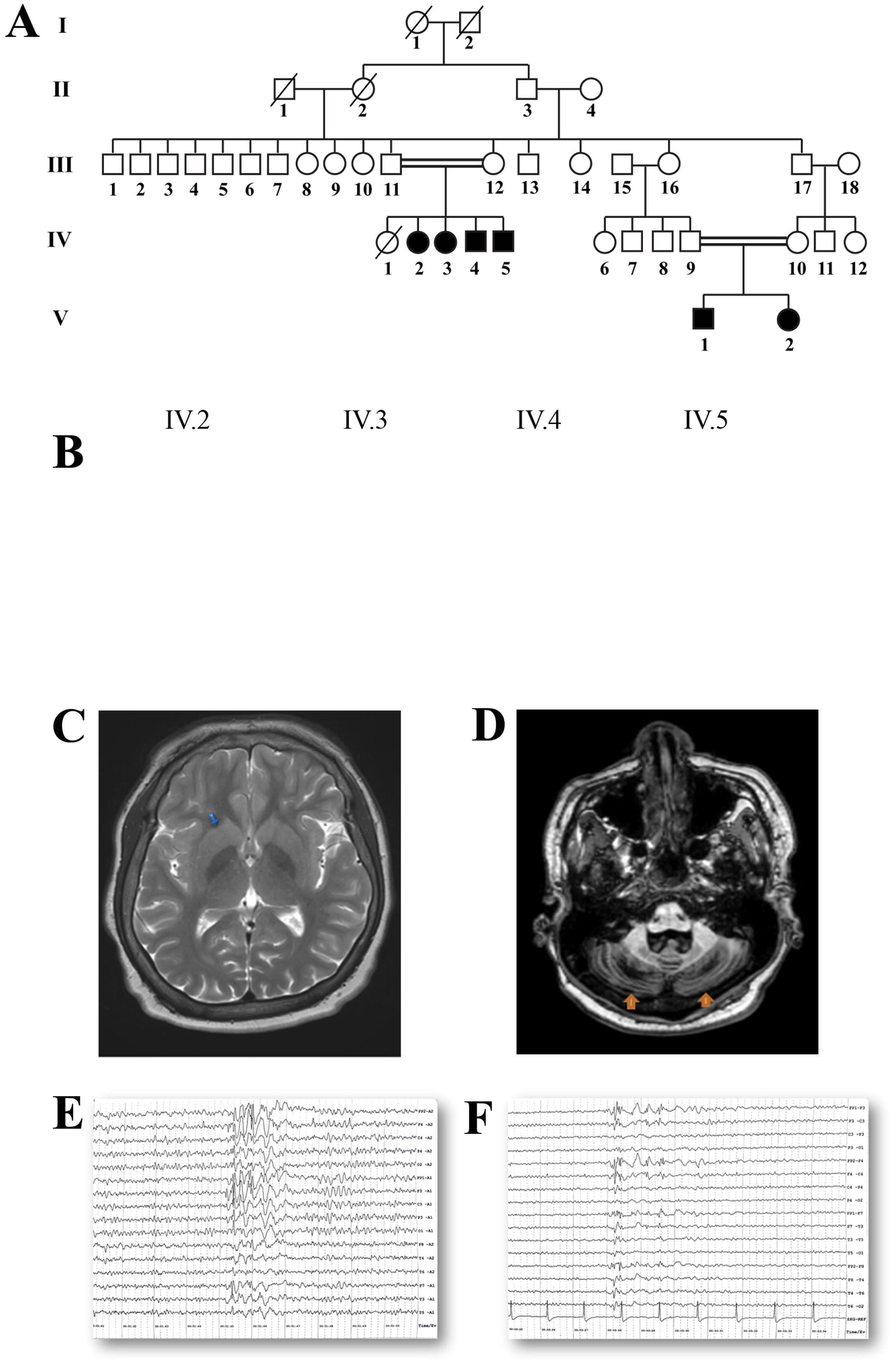
Clinical diagnosis of SeSAME family members. (A) Genogram of family with SeSAME syndrome with no electrolyte imbalance. The generations are marked in roman letters (I to V) and individuals in each generation are given running numbers. (B) All affected siblings showed dysmorphic facial features. (C) T2W image of IV.2 showing enlarged and bilateral basal ganglia (blue arrows) (D) T1 MPRAGE of IV.2 showing bilateral cerebellar atrophy (orange arrows) (E) EEG of V.1 showing generalized sharp and slow wave discharges predominantly in Fronto Central region (F) EEG of V.2 showing generalized poly spike discharges predominantly in Fronto -Temporal region.

### Variant prioritization using ROH and non-ROH methods identified two novel variants in *KCNJ10* and *PI4KB* and revealed mutation burden in Chr 1 in all patients

To identify the critical disease-associated loci, we performed WES and prioritized variants based on two independent approaches; assessing the exome-wide levels of homozygosity (ROH method) and assessing variants based on allele frequencies with autosomal recessive inheritance pattern (non-ROH method) in all family members. Unanimously, both analysis pipelines identified two novel high-risk disease-associated variants in *KCNJ10* and *PI4KB* and five rare variants in *PVRL4, RORC, FLG2, FCRL1*, and *NIT1* and one common variant in *HSPA6* segregating in homozygous state in all patients and heterozygous state in both parents. Surprisingly, both methods revealed mutational burden in Chr1 (Fig. 2a; Table 2).

**Figure 2.**
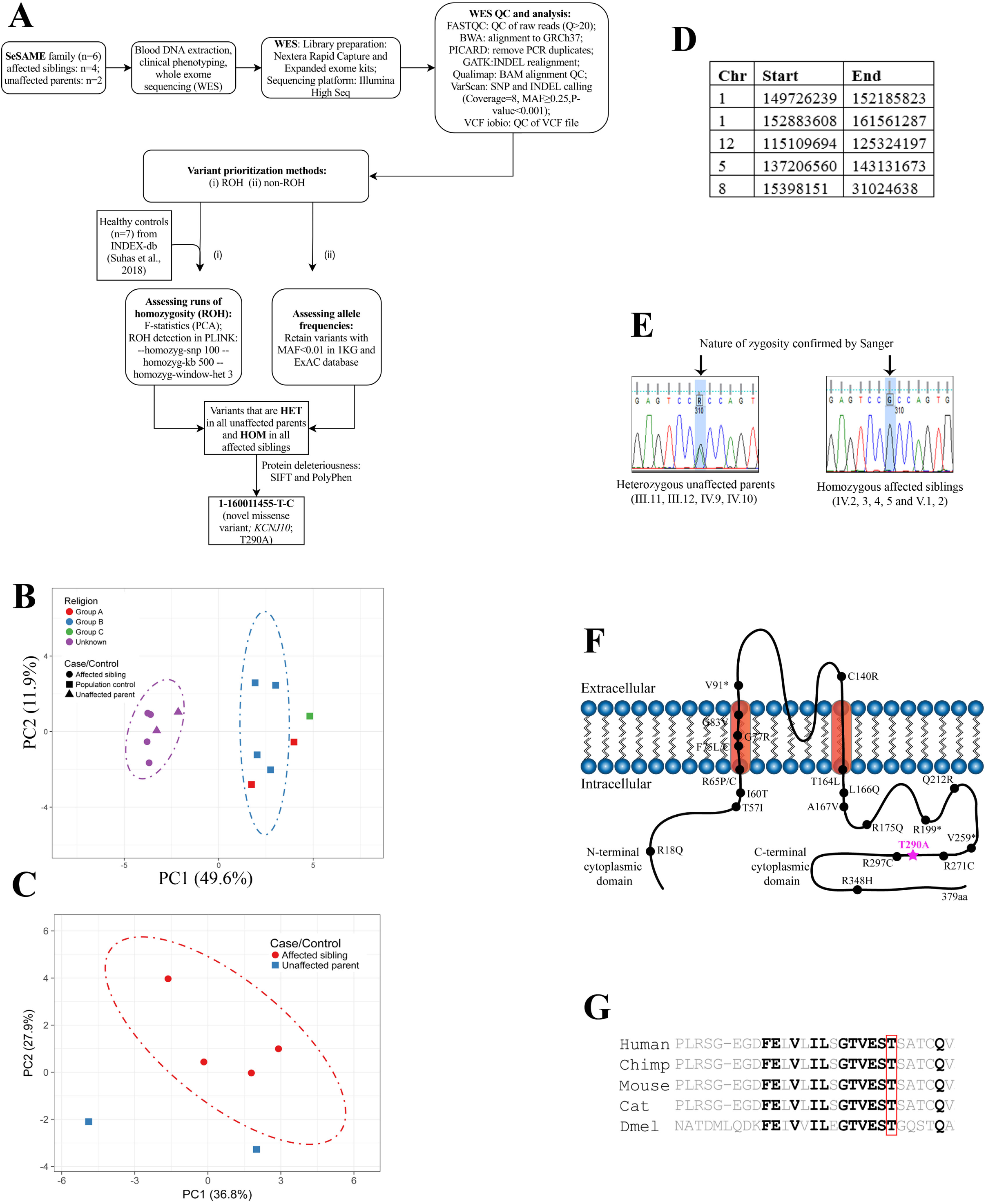
Identification of novel mutation in *KCNJ10* by homozygosity mapping and whole exome analysis of SeSAME family members. (A) WES analysis pipeline and variant prioritization methods. (B) Principle component analysis (PCA) of exome-wide F-statistics explains for an overall variance of ∼49% (PC1) between the SeSAME family members (purple ellipse) and healthy population controls (blue ellipse). The dot-dash lines in the plot represents the 95% confidence ellipse. (C)PCA plot explaining intra-familial levels of homozygosity between affected and un-affected members. (D) ROH regions observed in all patients but not in parental controls. (E) The zygosity of the *KCNJ10*^T290A^ variant was validated in all the six affected (HOM) and the four unaffected individuals (HET) within the pedigree. (F) A schematic reconstruction of Kir4.1with the T290A variant (purple) mapped in the cytoplasmic C-terminal domain, along with other deleterious variants identified from previous studies. (G) Multiple sequence alignment (MSA) of the Kir4.1 protein sequence across species reveals the evolutionary conservation of T290A in VEST domain.

Deleterious genetic effects of inbreeding are evident in children born out of consanguineous unions with a relatively higher burden of homozygous alleles [34, 35, 36]. These effects have been implicated to influence the evolution of mental illness and neurodevelopmental disorders [34]. Since SeSAME syndrome follows autosomal recessive (AR) inheritance and the role of homozygous alleles in AR illness has been well established [37], we analyzed the exome-wide levels of homozygosity for all samples within the pedigree including seven population controls (see materials and methods). Principal Component Analysis (PCA) of the exome-wide F-statistics separates the family members (n=6) from the population controls (n=7), explaining for an overall variance of 49.6%. All samples (both familial and population) within the two clusters, fell within their 95% confidence ellipses, except for two controls representing the relatively admixed communities (Fig. 2b). The SeSAME family alone was subjected to PCA in which the cases (n=4) formed a cluster and the unaffected parents (n=2) fell outside the 95% confidence ellipse (Fig. 2c), explaining the intra-familial variance in homozygosity. The ROH within the exomes of the individuals in the pedigree and the population controls were identified. A total of 56 homozygous stretches (either overlapping or unique) were identified in all cases and controls, of which 44 stretches belonged to the four affected siblings and the remaining were distributed between unaffected parents and population controls (Supplementary Table 1). Nevertheless, no ROH was detected in a subset of population controls. The burden of ROHs witnessed in the cases as compared to controls could be attributed to their consanguineous parentage. Of the ROHs identified in total, five stretches were explicitly shared between all the affected siblings but not observed in the unaffected parents and population controls, which will henceforth be notified as ROH_affected_ (Fig. 2d). TheROH_affected_ consists of a union set of 5329 variants across all the cases and controls, of which any given variant was observed in at least one sample. Since the disorder follows an autosomal recessive (AR) inheritance pattern, of the 5329 variants, we identified those that were heterozygous (HET) in both unaffected parents, but homozygous (HOM) in all of the affected siblings. Seventy-eight such variants, belonging to 47 genes, were identified and all of them mapped to Chr 1 (Supplementary Table2). This skewed observation could not be attributed to the length of Chr 1 for three reasons: i) the method used to compute ROH uses a sliding window approach which essentially removes the bias induced by the length of the chromosome; ii) the same Chr 1 ROH was not observed in either of the controls; iii) no ROH was observed in Chr 2 despite its genomic length being comparable to that of Chr 1. Of the 78 variants only three missense variants i.e., i) Chr1:158368964-C-T (*OR10T2*) ii) Chr1:160011455-T-C (*KCNJ10*) and iii) Chr1:161495040-C-T (*HSPA6*), were predicted to be deleterious by two algorithms.

To identify other deleterious variants segregating within the family by AR pattern, which could have otherwise been ignored by the ROH based method, we identified all the exonic and splice variants (including non-synonymous, stop gain and stop loss). The common variants i.e., those with a minor allele frequency (MAF)>0.01 in 1KG_all (1000 Genomes Project) and ExAC_all (Exome Aggregation Consortium) databases, were excluded from the analysis. We identified seven variants belonging to seven genes (Supplementary Table3). Interestingly, all the seven variants were located within Chr1:151288779-161088292, which was a subset of ROH_affected_ (Fig. 2d). Among the seven variants, Chr1:160011455-T-C [*KCNJ10*] was an obvious overlap. The remaining six variants fell on *PI4KB, RORC, FLG2, FCRL1, PVRL4* and *NIT1* genes. Apart from *KCNJ10* variant, none were predicted to be deleterious by all six prediction algorithms. However, three of the remaining six variants (Chr1:151288779-T-C [*PI4KB*], Chr1:161049499-G-A [*PVRL4*] and Chr1:161088292-A-G [*NIT1*]) were predicted to be deleterious by at least two algorithms (Table 2). Finally, the zygosity of the *KCNJ10* variant was confirmed by sanger sequencing for six patients and four unaffected parents in the family (III.11-12, IV.2-5, IV.9-10 and V.1-2) (Fig. 2e).

Thus, of the union set of nine putative deleterious variants (three based on ROH method and seven based on allele frequencies) segregating within the family, the *KCNJ10* gene was shortlisted for functional analysis to unravel the molecular impact of the variant for following reasons: i) *KCNJ10*, the candidate gene known to cause SeSAME syndrome (Celmina et al., 2018); ii) the variant reported in the patients is novel; iii) this was the only deleterious variant identified by both methods and iv) the variant reside at the interface between transmembrane and cytoplasmic domain at the membrane (Fig. 2f) which is strongly conserved through evolution (Fig. 2g).

### Novel *KCNJ10* variant disrupts channel properties in patient-derived LCLs

LCLs have been routinely used as a surrogate *in vitro* cell model to investigate cellular mechanisms of neurodevelopmental psychiatric disorders [38]. To investigate the functional role of Kir4.1^T290A^, we generated patient-specific LCLs, validated by karyotype for six members of SeSAME family. All six LCLs are free from both numeric and structural chromosomal abnormalities (data not shown).

The barium-sensitive inwardly-rectifying K^+^ current in LCLs measured by whole-cell patch clamp was substantially compromised in all patients. Kir4.1^T290A^ significantly depolarized LCL membranes and showed deficits in clearance of extracellular K^+^. To determine whether LCLs express functionally active endogenous Kir4.1 protein, we used immunofluorescence (IF), western blot and electrophysiology (Fig. 3). In parental controls, Kir4.1 is in close proximity with the actin-rich plasma membrane, diffusely discernible in the cytoplasm and enriched in the nuclear membrane and nucleus (Fig. 3a). However, in all affected individuals, we observed an increased punctate distribution of Kir4.1 in the cytoplasm but with no apparent disparity in the nucleus and nuclear membrane (Fig. 3b). To confirm the IF findings, western blot analysis showed a substantial increase in the expression of Kir4.1 in all patients compared with unaffected parents (Fig. 3C and 3D). These findings suggest an unstable nature of the mutant Kir4.1^T290A^ in all patients.

**Figure 3.**
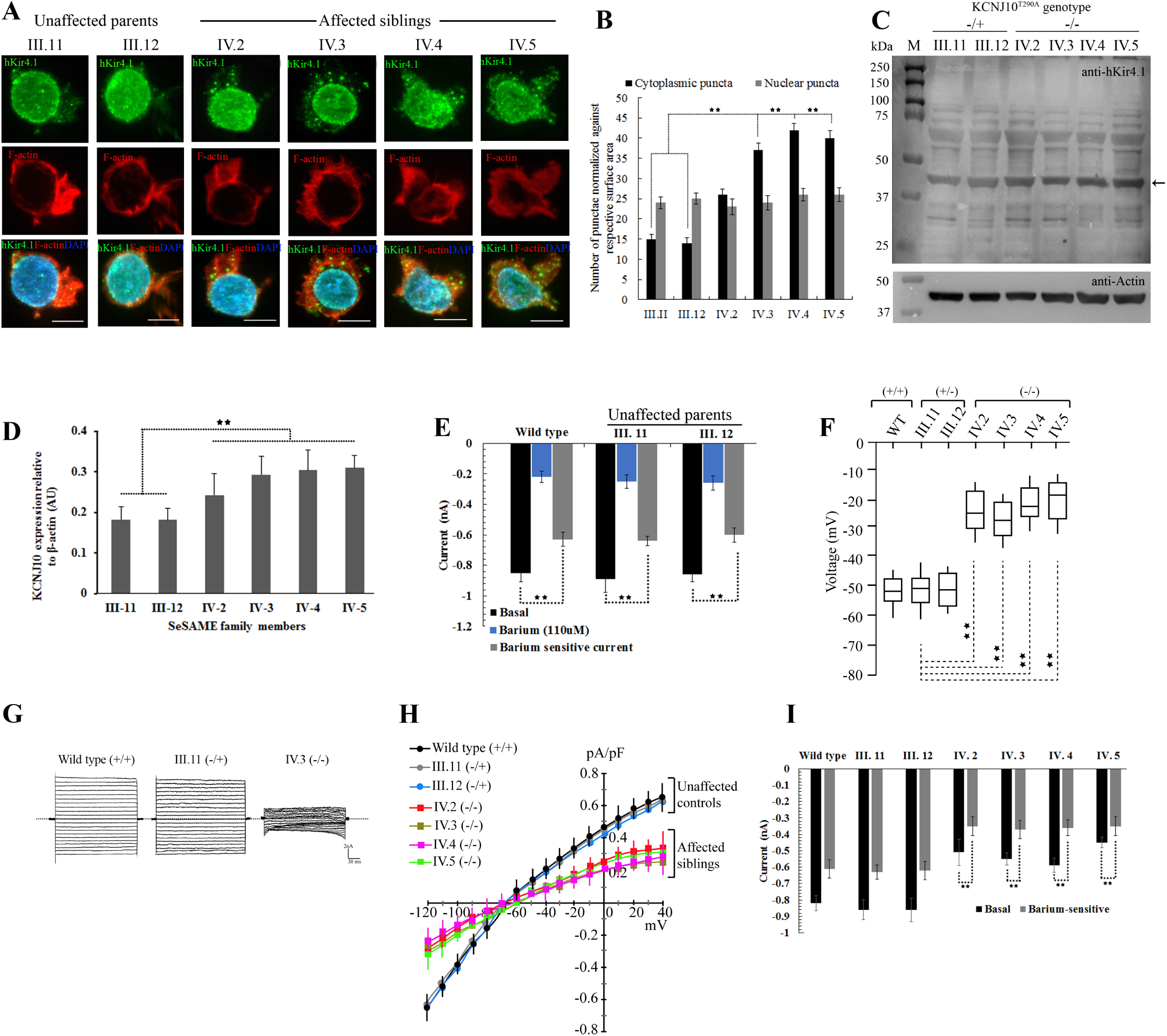
Novel Kir4.1^T290A^ mutation affects channel localization and function in patient-derived LCLs. (A) Projected *Z*-stacks of six LCLs showing the distribution of Kir4.1 in green, phalloidin to label F-actin in red and DAPI to label nucleus in blue. Scale bar, 10μm. (B) Quantitative measurement of cytoplasmic and nuclear punctae normalized against the cytoplasmic space (as measured by F-actin distribution) and nuclear space (as measured by DAPI distribution) in *Z*-stacks. (C) Anti-hKir4.1 western of six LCLs showing the distribution of both monomeric and multimeric forms of the protein. Arrow indicates the expression of Kir4.1 protein against beta-actin loading control (blot insert at the bottom). −/+ and −/− indicates the nature of zygosity of unaffected parents and affected individuals. (D) Densitometric plots representing the relative expression Kir4.1 protein from three independent western experiments is represented as mean±SE. Data analyzed using ANOVA. (E) Whole-cell currents measured from healthy wild type controls and two unaffected parental controls in response to voltage step protocol from −120 to 40mV in presence and absence of 110μM barium. Cells were clamped at *Vm*, equal to resting *Vm (Vh=Vm)*. Histogram shows the subtraction of currents obtained with barium from whole-cell currents, which served as internal control for each experiment. Barium sensitive current shows the contribution of Kir channels to whole-cell currents in each LCLs. Data analysed by *k* independent Kruskal-Wallis test with Bonferroni correction and represented as ±S.E. (F) Average membrane potential of LCLs from healthy control (wild type), two unaffected parents (III.11 and III. 12) and four affected (IV.2 to IV.4). Data analyzed using *k* independent group one-way ANOVA test with Turkey-Kramer post hoc tests. (G) whole-cell patch clamp recordings in response to voltage-steps from −120 to 40mV in 10mV steps, from a holding potential of −30mV. Representative currents traces from respective LCLs. (H) Current-voltage relationship is summarized within −120 to 40mV range. (I) Summary of inward currents discharges measured in response to induced K^+^ steps from 5-20 mM extracellular K^+^. For improved Kir specificity, Kir current discharges measured with and without barium. Data analysed using *k* independent group one-way ANOVA test with Turkey-Kramer post hoc tests. Error bars represent ±S.E. ** represents *p*<0.001

To confirm whether the endogenous Kir4.1 expressed in LCLs is functionally active and elicit detectable inward-rectifying potassium currents *in vitro*, we performed whole-cell patch clamp recordings in response to voltage-steps from −120 to 40mV in 10mV, from a holding potential of − 30mV both in the presence and absence of 110μM barium, a selective Kir channel blocker. Baseline current discharges from two heterozygous parental controls (III.11: −0.89±0.086, n=18, *p*=1.114 and III.12: −0.86±0.049, n=16, *p*=1.347) were not significantly different from wild type controls (−0.85±0.046, n=17) (Fig. 3E). In contrast, the average barium-sensitive current densities were substantially decreased in all three control LCLs tested, in heterozygous parents (III.11: − 0.64±0.041, n=15, *p*=2.1E-4 and III.12: −0.60±0.086, n=14, *p*=1.8E-4) and wild type (−0.63±0.104, n=14, *p*=2.5E-4) compared with their respective baseline discharges, implying the specificity of K^+^ currents recorded from endogenous Kir channels (Fig. 3E).

We recorded the resting membrane potential of LCLs from patients (Fig. 3F). Average membranes voltages from all patients (IV.2: −30mV±3.640, n=18, *p*=1.3E-5; IV.3: −32mV±2.156, n=20, *p*=2.4E-5; IV.4: −31mV±3.083, n=17, *p*=1.7E-4; IV.5: −24mV±2.817, n=20, *p*=2.8E-5) were significantly hyperpolarized as compared to wild type (WT:-55mV±4.102, n= 24) and parental controls (III.11: −51mV±3.842, n=21 and III.12: −50mV±4.21, n=19). In whole-cell voltage clamp, membrane current amplitudes were measured in all family members at both positive and negative potentials than the K^+^ equilibrium potential (E_k_) (Fig. 3G and 3H). The mean current densities as a function of voltage (pA/pF) measured in all those expressing the mutant channel were markedly smaller than wild type and parental controls (Fig. 3H). One major facet of the Kir4.1 channel is to clear extracellular K^+^ thereby showing stronger rectification. To test the K^+^ clearance ability of LCLs, we clamped the cells at their resting membrane potential, with and without 110μM barium, and measured the elicited membrane current discharges upon induced K^+^steps (from 5-20 mM). Overall, barium-sensitive currents from all patients were significantly reduced when compared to both parental and wild type controls (Fig. 3I).

## Discussion

In this study, we identified two novel pathogenic variants in *KCNJ10* and *PI4KB*, five rare pathogenic variants in *PVRL4*, *RORC*, *FLG2, FCRL1* and *NIT 1* and one common pathogenic variant in *HSPA6* suggesting the importance of membrane lipid signaling, adhesion-mediated cell migration and protein trafficking in SeSAME syndrome through regulation of Kir channel activity. In multiple biological contexts, these cellular processes are tightly linked in regulating Kir4.1 channel function at the plasma membrane [9, 39, 40, 41, 42, 43]. Functional studies in patient-specific LCLs suggests that the variant in *KCNJ10* causes 60% reduction in Kir4.1 channel activity which is presumably due to altered protein localization and decreased surface expression of mutant proteins. Finally, our study identified risk-associated variants in seven new genes in SeSAME syndrome, which might act as modifiers by regulating Kir4.1 channel function. A detailed mechanistic study investigating the biology of these modifiers in Kir4.1 physiology will help us to underpin the biology of disease manifestation in SeSAME syndrome.

Signal-dependent Golgi export processes have been implicated in Andersen-Tawil syndrome (ATS1) by controlling the surface density Kir2.1 channel [44]. It has become evident in recent years, that differential trafficking of Kir channels controls neuronal excitability, hormone secretion, action potential, K^+^ homeostasis and salt balance. The shared Golgi export signal patch at the cytoplasmic region in Kir2.3 and Kir4.1 is an AP-1 clathrin adaptor recognition site which ensures an additional quality control check point for the exit of mature folded channels [39]. The variant reported in this study Kir4.1^T290A^, reside in close proximity to Golgi export patch at the cytoplasmic region, implying the role of protein trafficking in SeSAME syndrome. Supporting this view, non-ROH method of analysis identified a pathogenic common variant in *HSPA6* gene, a molecular chaperone involved in ATP-dependent protein quality control system. It is also interesting to note the association of *HSPA6* variant in patients with sensory disturbances [45] suggesting mutations in genes that regulate protein trafficking can influence surface expression of Kir4.1 channel, irrespective of its variants.

All six patients reported here displayed relatively uniform and expected neurological and psychiatric manifestations, but they did not manifest electrolyte imbalance. Therefore, how and why certain *KCNJ10* variants fail to manifest electrolyte imbalance in SeSAME syndrome needs to be explored. There could be two possibilities for this discrepancy. First, it’s possible that certain *KCNJ10* mutations can affect CNS functions independently of other organ systems. It is conceivable that astrocytes and microglial cells of nervous system are highly sensitive to dysregulation of potassium homeostasis, while basolateral membrane in the distal nephron may be impervious to this effect [26]. Another possibility is that same *KCNJ10* variants could behave differently between CNS and kidney, since the channel activity depends largely on the formation of heterotetramers with other Kir entities (Kir5.1), cell type specificity, gating mechanisms and its influence on cell surface signaling receptors through PIP2 binding [9, 40, 43, 46]. In addition, it is unclear whether renal electrolyte deficit is a progressive impairment that develops over time, or a direct effect of the mutation, which necessitates further investigations and follow-up clinical evaluations. These different mechanisms suggest that although major gene effects are probably the primary drivers of illness, the diversity in clinical presentation is perhaps an outcome of complex genetic interactions between common and rare variants, each of varying effect sizes.

Surprisingly, both methods concluded a mutational and ROH burden in Chr 1. Given the clinical diversity and for additional reasons as discussed above, we suggest two possibilities for ROH and mutational burden which are broadly classified into intrinsic and extrinsic factors. Intrinsic factors include recombination hot-spots, defects in DNA repair, chromatin remodelling and yet unidentified intra-cellular signaling events, that favour to the occurrence of ROH, co-segregating with the illness. The extrinsic factor could be the clan structure of the family, which indicates a high-degree of endogamy. Another possibility is that individual ROHs might play key role in spatial-temporal regulation of gene expression within cell types that are sensitive to K^+^ homeostasis. The difference in the expression of Kir4.1 in patients in our SeSAME pedigree also highlights the role of ROH in gene regulation. Therefore, it would be helpful to investigate the functional consequences of homozygosity in expression of genes within the ROH and/or in close proximity especially in cell types that are relevant to the pathophysiology of SeSAME syndrome. Finally, an interplay between these factors could help us discriminate the cause and effect relationship of ROH in clinical diversity of SeSAME syndrome. Usually for every pregnancy in autosomal recessive disorders, there is a probability of 0.25 that the offspring(s) will inherit two copies of the disease gene and will therefore exhibit the phenotype [47]. However, in a clinical setting this distribution is skewed more towards almost all affected individuals in the same generation, than one would rather expect by chance, especially in children born to consanguineous unions. Thus, this skewed observation needs to be addressed at holistic paradigms by developing bio-physical and mathematical models to understand the physics and governing dynamics of the intra-cellular events, influencing the silent recombination choices of homologous chromosomes.

Though our study identified novel and common variants in new genes and its pathways that could help modify the activity of Kir channels in SeSAME pathogenesis, a complete mechanistic understanding would require establishment of animal models to explore the cell-type specific role of Kir4.1 in brain function. Justifying the importance of K^+^ homeostasis in brain, Kir4.1 knockout mouse, *Xenopus*, *zebrafish* and *Drosophila* mimics a subset of SeSAME symptoms in humans [6, 7, 10, 26, 29, 48, 49]. Therefore, future experiments with *in vivo* model systems will help dissect the cross talk of Kir4.1 signaling with membrane lipids [50], cell adhesion in axon guidance and synaptic architecture which is an essential feature for proper synaptic transmission and plasticity.

Our study identified two novel and five rare variants in genes that potentially modifies the channel properties of Kir4.1-mediated pathogenesis in SeSAME syndrome. In future, genetic interaction experiments in cell and/or animal model systems will help us tease apart the causative effects of these novel modifiers in Kir4.1 biology.

## Acknowledgement

The authors are immensely grateful to all members of SeSAME family for their participation and constant involvement in this study. We thank Dr. Gautham Arunachal U (NIMHANS) for describing the various dysmorphic features of the SeSAME kindreds. We thank ADBS genomic team for sharing the WES analysis pipeline. This work was generously supported with funds from Ramalingaswami re-entry fellowship (RLF/DBT/2015), ADBS (BT/PR17316/MED/31/326/2015) from Department of Biotechnology and Department of Science and Technology (ECR/2015/000468). Finally, the authors are thankful to all members of the molecular genetics, ADBS lab and the ADBS consortium for suggestions and discussions during the course of investigation.

## Funding

These experiments were supported by Ramalingaswami re-entry fellowship (RLF/DBT/2015) and ADBS (BT/PR17316/MED/31/326/2015) from Department of Biotechnology; Early career grant (ECR/2015/000468) from Department of Science and Technology.

## Competing interests

The authors declare no competing or financial interests.

## References

1 Klassen T, Davis C, Goldman A, Burgess D, Chen T, Wheeler D, McPherson J, Bourquin T, Lewis L, Villasana D, Morgan M, Muzny D, Gibbs R, Noebels. Exome sequencing of ion channel genes reveals complex profiles confounding personal risk assessment in epilepsy. J.Cell. 2011 Jun 24;145(7):1036–48.

2 Imbrici P, Camerino DC, Tricarico D. 2013. Major channels involved in neuropsychiatric disorders and therapeutic perspectives. Front Genet. 2013 May 7;4:76.

3 Begum R, Bakiri Y, Volynski KE, Kullmann DM. Action potential broadening in a presynaptic channelopathy. Nat Commun. 2016 Jul 6;7:12102.

4 Noebels J. Precision physiology and rescue of brain ion channel disorders.J Gen Physiol. 2017 May 1;149(5):533–546.

5 Middleton SJ, Kneller EM, Chen S, Ogiwara I, Montal M, Yamakawa K, McHugh TJ.Altered hippocampal replay is associated with memory impairment in mice heterozygous for the Scn2a gene. Nat Neurosci. 2018 Jul;21(7):996–1003.

6 Schirmer L, Möbius W, Zhao C, et al. Oligodendrocyte-encoded Kir4.1 function is required for axonal integrity. Elife. 2018;7:e36428.

7 Larson VA, Mironova Y, Vanderpool KG, Waisman A, Rash JE, Agarwal A, Bergles DE. Oligodendrocytes control potassium accumulation in white matter and seizure susceptibility. eLife. 2018;7:e34829.

8 Ye M, Yang J, Tian C, Zhu Q, Yin L, Jiang S, Yang M, Shu Y. Differential roles of NaV1.2 and NaV1.6 in regulating neuronal excitability at febrile temperature and distinct contributions to febrile seizures.Sci Rep. 2018 Jan 15;8(1):753.

9 deHart GW, Jin T, McCloskey DE, Pegg AE, Sheppard D. The alpha9beta1 integrin enhances cell migration by polyamine-mediated modulation of an inward-rectifier potassium channel. Proc Natl Acad Sci U S A. 2008;105(20):7188–93.

10 Kelley KW, Ben Haim L, Schirmer L, et al. Kir4.1-Dependent Astrocyte-Fast Motor Neuron Interactions Are Required for Peak Strength. Neuron. 2018;98(2):306–319.e7.

11 Lupski JR, Belmont JW, Boerwinkle E, Gibbs RA.Clan genomics and the complex architecture of human disease.Cell. 2011 Sep 30;147(1):32–43.

12 Bockenhauer, D., Feather, S., Stanescu, H. C., Bandulik, S., Zdebik, A. A., Reichold, M., et al. Epilepsy, ataxia, sensorineural deafness, tubulopathy, and KCNJ10 mutations. N. Engl. J. Med. 2009; 360: 1960–1970.

13 Scholl UI, Choi M, Liu T, et al. Seizures, sensorineural deafness, ataxia, mental retardation, and electrolyte imbalance (SeSAME syndrome) caused by mutations in KCNJ10. Proc Natl Acad Sci U S A. 2009;106(14):5842–7.

14 Celmina M, Micule I, Inashkina I, Audere M, Kuske S, Pereca J, Stavusis J, Pelnena D, Strautmanis J. EAST/SeSAME syndrome: Review of the literature and introduction of four new Latvian patients. Clin Genet. 2018 May 3.

15 Heuser K, Nagelhus EA, Taubøll E, Indahl U, Berg PR, Lien S, Nakken S, Gjerstad L, Ottersen OP. Variants of the genes encoding AQP4 and Kir4.1 are associated with subgroups of patients with temporal lobe epilepsy. Epilepsy Research 2010; 88:55–64.

16 Sicca F, Imbrici P, D’Adamo MC, Moro F, Bonatti F, Brovedani P, et al. Autism with seizures and intellectual disability: possible causative role of gain-of-function of the inwardly-rectifying K+ channel Kir4.1. Neurobiology of Disease 2011; 43:239–247.

17 Sicca F, Ambrosini E, Marchese M, Sforna L, Servettini I, Valvo G, et al. Gain-of-function defects of astrocytic Kir4.1 channel in children with autism spectrum disorders and epilepsy. Scientific Reports 2016; 6:34325.

18 Tong X, Ao Y, Faas GC, Nwaobi SE, Xu J, Haustein MD, et al. Astrocyte Kir4.1 ion channel deficit contributes to neuronal dysfunction in Huntington’s disease model mice. Nat. Neurosci. 2014; 17, 694–703.

19 Gu C. KIR4.1: K+ Channel Illusion or Reality in the Autoimmune Pathogenesis of Multiple Sclerosis. Front Mol Neurosci. 2016;9:90.

20 Kahanovitch U, Cuddapah VA, Pacheco NL, Holt LM, Mulkey DK, Percy AK, Olsen ML. MeCP2 Deficiency Leads to Loss of Glial Kir4.1. eNeuro. 2018: 19; 5(1). 0194–17.

21 Martin HC, Jones WD, McIntyre R, Sanchez-Andrade G, Sanderson M, Stephenson JD et al. A SINE Insertion in ATP1B2 in Belgian Shepherd Dogs Affected by Spongy Degeneration with Cerebellar Ataxia (SDCA2). G3 (Bethesda). 2017;7(8):2729–2737.

22 Van Poucke M, Stee K, Bhatti SF, et al. The novel homozygous KCNJ10 c.986T>C (p.(Leu329Pro)) variant is pathogenic for the SeSAME/EAST homologue in Malinois dogs. Eur J Hum Genet. 2016; 25(2):222–226.

23 Lopatin AN, Makhina EN, Nichols CG. Potassium channel block by cytoplasmic polyamines as the mechanism of intrinsic rectification. Nature1994; 24:372(6504):366–9.

24 Paulais M, et al. Renal phenotype in mice lacking the Kir5.1 (Kcnj16) K+ channel subunit contrasts with that observed in SeSAME/EAST syndrome. Proc Natl Acad Sci USA. 2011;108(25):10361–10366.

25 Kofuji P, Ceelen P, Zahs KR, Surbeck LW, Lester HA, Newman EA. Genetic inactivation of an inwardly rectifying potassium channel (Kir4.1 subunit) in mice: phenotypic impact in retina. J Neurosci. 2000;20:5733–5740.

26 Djukic B, Casper KB, Philpot BD, Chin LS, McCarthy KD. Conditional knock-out of Kir4.1 lead to glial membrane depolarization, inhibition of potassium and glutamate uptake, and enhanced short-term synaptic potentiation. J Neurosci. 2007;27:11354–11365.

27 Neusch C., Papadopoulos N., Muller M. et al. Lack of the Kir4.1 channel subunit abolishes K+buffering properties of astrocytes in the ventral respiratory group: impact on extracellular K+ regulation. J. Neurophysiol. 2006; 95, 1843–1852.

28 Song F, Hong X, Cao J, et al. Kir4.1 channel in NG2-glia play a role in development, potassium signaling, and ischemia-related myelin loss. Commun Biol. 2018;1:80.

29 Neusch C., Rozengurt N., Jacobs R. E., Lester H. A. and Kofuji P. Kir4.1 potassium channel subunit is crucial for oligodendrocyte development and in vivo myelination. J. Neurosci. 2001; 21, 5429–5438

30 Hui-Yuen J, McAllister S, Koganti S, Hill E, Bhaduri-McIntosh S. Establishment of Epstein-Barr virus growth-transformed lymphoblastoid cell lines. Journal of visualized experiments: JoVE. 2011(57).

31 Suhas G, Husayn Ahmed P, Ravi Kumar Nadella, Ravi Prabhakar More, Manasa Seshadri, Biju Viswanath, Mahendra Rao, Sanjeev Jain, The ADBS consortium, Odity Mukherjee. Exome sequencing in families with severe mental illness identifies novel and rare variants in genes implicated in Mendelian neuropsychiatric syndromes. Psychiatry and Clinical Neurosciences 2018.

32 Ahmed P, Vidhya V, et al. INDEX-db: The Indian Exome Reference database (Phase-I). biorxiv. 2018.doi: https://doi.org/10.1101/312090

33 Purcell S, Neale B, Todd-Brown K, et al. PLINK: a tool set for whole-genome association and population-based linkage analyses. Am J Hum Genet. 2007;81(3):559–75.

34 Bittles AH, Black ML. Evolution in health and medicine Sackler colloquium: Consanguinity, human evolution, and complex diseases. Proc Natl Acad Sci U S A. 2010; 26:107.

35 Shawky RM, Elsayed SM, Zaki ME, et al. Consanguinity and its relevant to clinical genetics. Egyptian Journal of Medical Human Genetics. 2013;14:157–64.

36 Corry PC. Consanguinity and prevalence patterns of inherited disease in the UK Pakistani community. Hum Hered. 2014; 77(1-4): 207–16.

37 Sund KL, Zimmerman SL, Thomas C, Mitchell AL, Prada CE, Grote L, Bao L, Martin LJ, Smolarek TA. Regions of homozygosity identified by SNP microarray analysis aid in the diagnosis of autosomal recessive disease and incidentally detect parental blood relationships. Genet Med. 2013; 15(1):70–8.

38 Kumar S, Curran JE, Glahn DC, Blangero J. Utility of Lymphoblastoid Cell Lines for Induced Pluripotent Stem Cell Generation. Stem Cells Int. 2016; 2349261.

39 Li X, Ortega B, Kim B, Welling PA. A Common Signal Patch Drives AP-1 Protein-dependent Golgi Export of Inwardly Rectifying Potassium Channels. J Biol Chem. 2016 Jul 15;291(29):14963–72.

40 Hansen SB, Tao X, MacKinnon R. Structural basis of PIP2 activation of the classical inward rectifier K+ channel Kir2.2. Nature. 2011 Aug 28;477(7365):495–8.

41 Ma D, Taneja TK, Hagen BM, Kim BY, Ortega B, Lederer WJ, Welling PA.Golgi export of the Kir2.1 channel is driven by a trafficking signal located within its tertiary structure.Cell. 2011 Jun 24;145(7):1102–15.

42 Du X, Zhang H, Lopes C, Mirshahi T, Rohacs T, Logothetis DE. Characteristic interactions with phosphatidylinositol 4,5-bisphosphate determine regulation of kir channels by diverse modulators. J Biol Chem. 2004 Sep 3;279(36):37271–81.

43 Lopes CM, Zhang H, Rohacs T, Jin T, Yang J, Logothetis DE. Alterations in conserved Kir channel-PIP2 interactions underlie channelopathies. Neuron. 2002 Jun 13;34(6):933–44.

44 Plaster, N.M., Tawil, R., Tristani-Firouzi, M., Canún, S., Bendahhou, S., Tsunoda, A., Donaldson, M.R., Iannaccone, S.T., Brunt, E., Barohn, R., et al. (2001). Mutations in Kir2.1 cause the developmental and episodic electrical phenotypes of Andersen’s syndrome. Cell 105, 511–519.

45 Kobayashi D, Nishizawa D, Takasaki Y, et al. Genome-wide association study of sensory disturbances in the inferior alveolar nerve after bilateral sagittal split ramus osteotomy. Mol Pain. 2013;9:34.

46 Reichold M, Zdebik AA, Lieberer E, et al. KCNJ10 gene mutations causing EAST syndrome (epilepsy, ataxia, sensorineural deafness, and tubulopathy) disrupt channel function. Proc Natl Acad Sci U S A. 2010;107(32):14490–5.

47 Ellard S, Kivuva E, Turnpenny P, et al. An exome sequencing strategy to diagnose lethal autosomal recessive disorders. Eur J Hum Genet. 2014;23(3):401–4.

48 Chen R, Swale DR. Inwardly Rectifying Potassium (Kir) Channels Represent a Critical Ion Conductance Pathway in the Nervous Systems of Insects. Sci Rep. 2018;8(1):1617.

49 Dahal GR, Pradhan SJ, Bates EA. Inwardly rectifying potassium channels influence Drosophila wing morphogenesis by regulating Dpp release. Development. 2017 Aug 1;144(15):2771–2783.

50 Hardie, R. C., Gu, Y., Martin, F., Sweeney, S. T. and Raghu, P. (2004). In vivo light induced and basal phospholipase C activity in Drosophila photoreceptors measured with genetically targeted phosphatidylinositol 4,5-bisphosphatesensitive ion channels (Kir2.1). J. Biol. Chem. 279, 47773–47782.

51 Ahmad F, Nasir A, Thiele H, Umair M, Borck G, Ahmad W. A novel homozygous missense variant in NECTIN4 (PVRL4) causing ectodermal dysplasia cutaneous syndactyly syndrome. Ann Hum Genet. 2018 Jul;82(4):232–238

52 Okada, S., Markle, J. G., Deenick, E. K., Mele, F., Averbuch, D., Lagos, M., Alzahrani, M et al Impairment of immunity to Candida and Mycobacterium in humans with bi-allelic RORC mutations. Science 349: 606–613, 2015.

