## Supplemental Table 1 for "Identification and functional characterization of two novel mutations in *KCNJ10* and *PI4KB* in SeSAME syndrome without electrolyte imbalance"

**Supplementary Table 1: ROH (n=56) (either overlapping or unique) detected within th**

| FID | IID | PHE | CHR | SNP1 | SNP2 | POS1 |
|---|---|---|---|---|---|---|
| Control | 10 | -9 | 11 | 0 | 0 | 207275 |
| Control | 10 | -9 | 11 | 0 | 0 | 1253980 |
| Control | 10 | -9 | 11 | 0 | 0 | 4510285 |
| Control | 10 | -9 | 17 | 0 | 0 | 76568872 |
| Control | 11 | -9 | 12 | 0 | 0 | 6132098 |
| Control | 11 | -9 | 12 | 0 | 0 | 52697806 |
| Control | 12 | -9 | 13 | 0 | 0 | 19775525 |
| Control | 13 | -9 | 23 | 0 | 0 | 149114513 |
| SeSAME | A1 | -9 | 11 | 0 | 0 | 5566030 |
| SeSAME | A1 | -9 | 22 | 0 | 0 | 39631547 |
| SeSAME | B2 | -9 | 1 | 0 | 0 | 31188089 |
| SeSAME | B2 | -9 | 9 | 0 | 0 | 129265612 |
| SeSAME | C3 | -9 | 1 | 0 | 0 | 149726239 |
| SeSAME | C3 | -9 | 1 | 0 | 0 | 152883608 |
| SeSAME | C3 | -9 | 2 | 0 | 0 | 219941063 |
| SeSAME | C3 | -9 | 2 | 0 | 0 | 227871797 |
| SeSAME | C3 | -9 | 5 | 0 | 0 | 137206560 |
| SeSAME | C3 | -9 | 8 | 0 | 0 | 15398151 |
| SeSAME | C3 | -9 | 11 | 0 | 0 | 43284469 |
| SeSAME | C3 | -9 | 12 | 0 | 0 | 94533886 |
| SeSAME | C3 | -9 | 22 | 0 | 0 | 49662324 |
| SeSAME | D4 | -9 | 1 | 0 | 0 | 149676075 |
| SeSAME | D4 | -9 | 1 | 0 | 0 | 152384833 |
| SeSAME | D4 | -9 | 2 | 0 | 0 | 131935190 |
| SeSAME | D4 | -9 | 4 | 0 | 0 | 4763704 |
| SeSAME | D4 | -9 | 8 | 0 | 0 | 15398151 |
| SeSAME | D4 | -9 | 12 | 0 | 0 | 115109694 |
| SeSAME | D4 | -9 | 12 | 0 | 0 | 130648299 |
| SeSAME | D4 | -9 | 13 | 0 | 0 | 19409651 |
| SeSAME | D4 | -9 | 15 | 0 | 0 | 96931594 |
| SeSAME | D4 | -9 | 16 | 0 | 0 | 2569810 |
| SeSAME | D4 | -9 | 16 | 0 | 0 | 86575992 |
| SeSAME | E5 | -9 | 1 | 0 | 0 | 109008064 |
| SeSAME | E5 | -9 | 1 | 0 | 0 | 149784642 |
| SeSAME | E5 | -9 | 1 | 0 | 0 | 152329369 |
| SeSAME | E5 | -9 | 3 | 0 | 0 | 7188116 |
| SeSAME | E5 | -9 | 3 | 0 | 0 | 182623765 |
| SeSAME | E5 | -9 | 5 | 0 | 0 | 130325939 |
| SeSAME | E5 | -9 | 10 | 0 | 0 | 42775352 |
| SeSAME | E5 | -9 | 10 | 0 | 0 | 70783075 |
| SeSAME | E5 | -9 | 10 | 0 | 0 | 87898729 |
| SeSAME | E5 | -9 | 10 | 0 | 0 | 111625108 |

| | | | | | | |
|---|---|---|---|---|---|---|
| SeSAME | E5 | -9 | 12 | 0 | 0 | 132834052 |
| SeSAME | E5 | -9 | 15 | 0 | 0 | 96931594 |
| SeSAME | E5 | -9 | 16 | 0 | 0 | 87782258 |
| SeSAME | E5 | -9 | 16 | 0 | 0 | 89299830 |
| SeSAME | E5 | -9 | 19 | 0 | 0 | 43257377 |
| SeSAME | F6 | -9 | 1 | 0 | 0 | 111731215 |
| SeSAME | F6 | -9 | 1 | 0 | 0 | 149726239 |
| SeSAME | F6 | -9 | 1 | 0 | 0 | 152883680 |
| SeSAME | F6 | -9 | 3 | 0 | 0 | 124748226 |
| SeSAME | F6 | -9 | 5 | 0 | 0 | 133328560 |
| SeSAME | F6 | -9 | 13 | 0 | 0 | 19042879 |
| SeSAME | F6 | -9 | 15 | 0 | 0 | 96931594 |
| SeSAME | F6 | -9 | 18 | 0 | 0 | 11619523 |
| SeSAME | F6 | -9 | 23 | 0 | 0 | 144904882 |

Supplementary Table 1

**ıe exomes of all cases and controls. No ROH detected in the populations controls 7, 8 an**

| POS2 | KB | NSNP | DENSITY | PHOM | PHET |
|---|---|---|---|---|---|
| 1010649 | 803.375 | 120 | 6.695 | 0.958 | 0.042 |
| 3602195 | 2348.216 | 106 | 22.153 | 0.981 | 0.019 |
| 11374283 | 6863.999 | 294 | 23.347 | 0.983 | 0.017 |
| 81169136 | 4600.265 | 189 | 24.34 | 0.963 | 0.037 |
| 7971833 | 1839.736 | 104 | 17.69 | 0.962 | 0.038 |
| 58378009 | 5680.204 | 236 | 24.069 | 0.975 | 0.025 |
| 21523002 | 1747.478 | 122 | 14.324 | 0.943 | 0.057 |
| 154456747 | 5342.235 | 121 | 44.151 | 0.959 | 0.041 |
| 6569896 | 1003.867 | 108 | 9.295 | 0.954 | 0.046 |
| 42523003 | 2891.457 | 107 | 27.023 | 0.944 | 0.056 |
| 35184033 | 3995.945 | 132 | 30.272 | 0.992 | 0.008 |
| 132377728 | 3112.117 | 176 | 17.682 | 0.977 | 0.023 |
| 152185823 | 2459.585 | 130 | 18.92 | 0.977 | 0.023 |
| 161561287 | 8677.68 | 480 | 18.079 | 0.992 | 0.008 |
| 220771378 | 830.316 | 123 | 6.751 | 0.992 | 0.008 |
| 234967539 | 7095.743 | 249 | 28.497 | 1 | 0 |
| 143131673 | 5925.114 | 194 | 30.542 | 0.974 | 0.026 |
| 33451026 | 18052.876 | 395 | 45.703 | 1 | 0 |
| 48346729 | 5062.261 | 148 | 34.204 | 0.993 | 0.007 |
| 125324197 | 30790.312 | 741 | 41.552 | 0.993 | 0.007 |
| 50893141 | 1230.818 | 153 | 8.045 | 0.98 | 0.02 |
| 152185972 | 2509.898 | 133 | 18.871 | 0.977 | 0.023 |
| 163309302 | 10924.47 | 547 | 19.972 | 0.989 | 0.011 |
| 132586953 | 651.764 | 135 | 4.828 | 0.963 | 0.037 |
| 8983355 | 4219.652 | 169 | 24.968 | 0.976 | 0.024 |
| 31024638 | 15626.488 | 371 | 42.12 | 1 | 0 |
| 129559557 | 14449.864 | 371 | 38.948 | 0.973 | 0.027 |
| 133808129 | 3159.831 | 177 | 17.852 | 0.966 | 0.034 |
| 23489920 | 4080.27 | 137 | 29.783 | 0.971 | 0.029 |
| 101922323 | 4990.73 | 123 | 40.575 | 0.984 | 0.016 |
| 11769963 | 9200.154 | 322 | 28.572 | 0.975 | 0.025 |
| 90233555 | 3657.564 | 323 | 11.324 | 0.969 | 0.031 |
| 117142531 | 8134.468 | 290 | 28.05 | 0.993 | 0.007 |
| 152186103 | 2401.462 | 143 | 16.793 | 0.986 | 0.014 |
| 161599693 | 9270.325 | 568 | 16.321 | 0.993 | 0.007 |
| 10885920 | 3697.805 | 131 | 28.228 | 0.969 | 0.031 |
| 188590650 | 5966.886 | 158 | 37.765 | 0.987 | 0.013 |
| 143200053 | 12874.115 | 328 | 39.25 | 0.985 | 0.015 |
| 46187373 | 3412.022 | 100 | 34.12 | 0.97 | 0.03 |
| 73537614 | 2754.54 | 160 | 17.216 | 0.981 | 0.019 |
| 106938110 | 19039.382 | 557 | 34.182 | 0.995 | 0.005 |
| 127584927 | 15959.82 | 401 | 39.8 | 0.985 | 0.015 |

| | | | | | |
|---|---|---|---|---|---|
| 133810703 | 976.652 | 111 | 8.799 | 0.973 | 0.027 |
| 102293008 | 5361.415 | 167 | 32.104 | 0.994 | 0.006 |
| 89299519 | 1517.262 | 208 | 7.295 | 0.986 | 0.014 |
| 90185214 | 885.385 | 129 | 6.863 | 0.969 | 0.031 |
| 43858317 | 600.941 | 119 | 5.05 | 0.958 | 0.042 |
| 117133188 | 5401.974 | 149 | 36.255 | 0.973 | 0.027 |
| 152186029 | 2459.791 | 139 | 17.696 | 0.986 | 0.014 |
| 161641237 | 8757.558 | 533 | 16.431 | 0.987 | 0.013 |
| 136310319 | 11562.094 | 303 | 38.159 | 0.993 | 0.007 |
| 143131673 | 9803.114 | 252 | 38.901 | 0.984 | 0.016 |
| 23490250 | 4447.372 | 173 | 25.707 | 0.977 | 0.023 |
| 102293008 | 5361.415 | 156 | 34.368 | 0.994 | 0.006 |
| 14183734 | 2564.212 | 104 | 24.656 | 0.971 | 0.029 |
| 155251726 | 10346.845 | 209 | 49.506 | 0.952 | 0.048 |

Supplementary Table 1
