## Supplemental table 2 for "Identification and functional characterization of two novel mutations in *KCNJ10* and *PI4KB* in SeSAME syndrome without electrolyte imbalance"

**Supplementary Table 2: Variants (n=78) within $ROH_{pro}$ that were HET in all of the unaffected**

| Chr-Pos-Ref-Alt | CHROM | POS | ID | REF | ALT | Control_7 |
|---|---|---|---|---|---|---|
| 1-152883680-A-G | 1 | 152883680 | . | A | G | 0/1 |
| 1-152883711-G-C | 1 | 152883711 | . | G | C | 0/1 |
| 1-152884111-G-A | 1 | 152884111 | . | G | A | 0/1 |
| 1-153012765-T-C | 1 | 153012765 | . | T | C | 0/1 |
| 1-153363105-T-C | 1 | 153363105 | . | T | C | 0 |
| 1-153588340-C-T | 1 | 153588340 | . | C | T | 0 |
| 1-154401972-T-C | 1 | 154401972 | . | T | C | 0 |
| 1-154426970-A-C | 1 | 154426970 | . | A | C | 0/1 |
| 1-154528053-G-A | 1 | 154528053 | . | G | A | 0/1 |
| 1-154544651-G-T | 1 | 154544651 | . | G | T | 0 |
| 1-154580458-C-A | 1 | 154580458 | . | C | A | 0 |
| 1-154899040-G-A | 1 | 154899040 | . | G | A | 0 |
| 1-155033317-T-C | 1 | 155033317 | . | T | C | 0 |
| 1-155034632-C-G | 1 | 155034632 | . | C | G | 0 |
| 1-155105882-T-G | 1 | 155105882 | . | T | G | 0 |
| 1-155106550-A-G | 1 | 155106550 | . | A | G | 0/1 |
| 1-155108287-G-A | 1 | 155108287 | . | G | A | 0 |
| 1-155142210-A-G | 1 | 155142210 | . | A | G | 0/1 |
| 1-155142229-G-A | 1 | 155142229 | . | G | A | 0/1 |
| 1-155148781-G-A | 1 | 155148781 | . | G | A | 0 |
| 1-155149718-G-A | 1 | 155149718 | . | G | A | 0/1 |
| 1-155162067-C-T | 1 | 155162067 | . | C | T | 0/1 |
| 1-156025096-C-T | 1 | 156025096 | . | C | T | 0 |
| 1-156202173-A-G | 1 | 156202173 | . | A | G | 0 |
| 1-156206158-C-T | 1 | 156206158 | . | C | T | 0 |
| 1-156213921-C-T | 1 | 156213921 | . | C | T | 0/1 |
| 1-156230206-C-A | 1 | 156230206 | . | C | A | 0 |
| 1-156261491-G-C | 1 | 156261491 | . | G | C | 0 |
| 1-156498673-T-A | 1 | 156498673 | . | T | A | 1/1 |
| 1-156499969-T-C | 1 | 156499969 | . | T | C | 1/1 |
| 1-156502870-C-T | 1 | 156502870 | . | C | T | 0 |
| 1-156641537-C-T | 1 | 156641537 | . | C | T | 0 |
| 1-156713558-G-A | 1 | 156713558 | . | G | A | 0 |
| 1-156761618-C-T | 1 | 156761618 | . | C | T | 0/1 |
| 1-156834364-G-A | 1 | 156834364 | . | G | A | 0 |
| 1-156882950-C-T | 1 | 156882950 | . | C | T | 0 |
| 1-156882996-G-C | 1 | 156882996 | . | G | C | 0 |
| 1-156883617-G-A | 1 | 156883617 | . | G | A | 0 |
| 1-157550066-T-C | 1 | 157550066 | . | T | C | 0 |
| 1-157768030-G-C | 1 | 157768030 | . | G | C | 0 |
| 1-157771880-C-T | 1 | 157771880 | . | C | T | 1/1 |
| 1-157802877-T-C | 1 | 157802877 | . | T | C | 0 |

| | | | | | | |
|---|---|---|---|---|---|---|
| 1-157802979-C-T | 1 | 157802979 | . | C | T | 0 |
| 1-157806006-G-A | 1 | 157806006 | . | G | A | 0 |
| 1-157894802-C-T | 1 | 157894802 | . | C | T | 0 |
| 1-158325891-T-C | 1 | 158325891 | . | T | C | 0 |
| 1-158368964-C-T | 1 | 158368964 | . | C | T | 0 |
| 1-158368974-A-G | 1 | 158368974 | . | A | G | 0 |
| 1-158369064-A-G | 1 | 158369064 | . | A | G | 0 |
| 1-158577167-A-T | 1 | 158577167 | . | A | T | 0 |
| 1-159505297-C-T | 1 | 159505297 | . | C | T | 0 |
| 1-159850401-C-G | 1 | 159850401 | . | C | G | 0 |
| 1-159897327-C-T | 1 | 159897327 | . | C | T | 0 |
| 1-159897754-C-T | 1 | 159897754 | . | C | T | 0 |
| 1-160011455-T-C | 1 | 160011455 | . | T | C | 0 |
| 1-160106531-G-A | 1 | 160106531 | . | G | A | 0 |
| 1-160134205-C-T | 1 | 160134205 | . | C | T | 0 |
| 1-160137300-A-C | 1 | 160137300 | . | A | C | 0 |
| 1-160146173-T-C | 1 | 160146173 | . | T | C | 0 |
| 1-160147502-T-G | 1 | 160147502 | . | T | G | 0 |
| 1-160147510-G-A | 1 | 160147510 | . | G | A | 0 |
| 1-161049499-G-A | 1 | 161049499 | . | G | A | 0 |
| 1-161088292-A-G | 1 | 161088292 | . | A | G | 0 |
| 1-161088583-T-G | 1 | 161088583 | . | T | G | 0 |
| 1-161092761-A-G | 1 | 161092761 | . | A | G | 0 |
| 1-161094190-A-G | 1 | 161094190 | . | A | G | 0 |
| 1-161126975-C-G | 1 | 161126975 | . | C | G | 0/1 |
| 1-161136843-C-G | 1 | 161136843 | . | C | G | 0 |
| 1-161139738-G-A | 1 | 161139738 | . | G | A | 0 |
| 1-161143801-G-A | 1 | 161143801 | . | G | A | 0 |
| 1-161145005-C-T | 1 | 161145005 | . | C | T | 0 |
| 1-161161284-G-C | 1 | 161161284 | . | G | C | 0 |
| 1-161172233-C-A | 1 | 161172233 | . | C | A | 0 |
| 1-161179877-G-A | 1 | 161179877 | . | G | A | 0 |
| 1-161495040-C-T | 1 | 161495040 | . | C | T | 0 |
| 1-161495117-C-T | 1 | 161495117 | . | C | T | 0 |
| 1-161495477-A-T | 1 | 161495477 | . | A | T | 0/1 |
| 1-161496134-C-G | 1 | 161496134 | . | C | G | 0 |

**l parents and HOM in all of the affected siblings. (0/1 = HET; 1/1 = HOM)**

| Control_8 | Control_9 | Control_10 | Control_11 | Control_12 | Control_13 | A1 |
|---|---|---|---|---|---|---|
| 0 | 0 | 0/1 | 0/1 | 0 | 0 | 0/1 |
| 0 | 0 | 0/1 | 0/1 | 0 | 0 | 0/1 |
| 0 | 0 | 0 | 0 | 0 | 0 | 0/1 |
| 0 | 0 | 0 | 0/1 | 0 | 0 | 0/1 |
| 0 | 0 | 0 | 0 | 0 | 0 | 0/1 |
| 0 | 0/1 | 0 | 0/1 | 0/1 | 0 | 0/1 |
| 0 | 0 | 0 | 0 | 0 | 0 | 0/1 |
| 0 | 0 | 0 | 0/1 | 0 | 0 | 0/1 |
| 0 | 0 | 0 | 0 | 0/1 | 0 | 0/1 |
| 0 | 0 | 0 | 0 | 0 | 0 | 0/1 |
| 0 | 0/1 | 1/1 | 0 | 0 | 0 | 0/1 |
| 0 | 0 | 0 | 0 | 0 | 0 | 0/1 |
| 1/1 | 1/1 | 0/1 | 1/1 | 1/1 | 0 | 0/1 |
| 0 | 0 | 0 | 0 | 0 | 0 | 0/1 |
| 0 | 0 | 0/1 | 0 | 0 | 0 | 0/1 |
| 1/1 | 1/1 | 0/1 | 0/1 | 0 | 0 | 0/1 |
| 0 | 1/1 | 0/1 | 0 | 0 | 0 | 0/1 |
| 1/1 | 0 | 0/1 | 0 | 0 | 0 | 0/1 |
| 1/1 | 0 | 0/1 | 0 | 0 | 0 | 0/1 |
| 0 | 0 | 0 | 0 | 0 | 0 | 0/1 |
| 0 | 0/1 | 0/1 | 0/1 | 1/1 | 0 | 0/1 |
| 0/1 | 0/1 | 0/1 | 1/1 | 1/1 | 0 | 0/1 |
| 0 | 0 | 0 | 0 | 0 | 0 | 0/1 |
| 0/1 | 1/1 | 0/1 | 0/1 | 0 | 0 | 0/1 |
| 0 | 0 | 0 | 0 | 0 | 0 | 0/1 |
| 0 | 0 | 0 | 0 | 0 | 0 | 0/1 |
| 0 | 0 | 0 | 0/1 | 0 | 0/1 | 0/1 |
| 0 | 0 | 0 | 0 | 0 | 0 | 0/1 |
| 0 | 0 | 0 | 0 | 0 | 0 | 0/1 |
| 1/1 | 0/1 | 1/1 | 1/1 | 1/1 | 0 | 0/1 |
| 0 | 0 | 0 | 0/1 | 0/1 | 0 | 0/1 |
| 0 | 0/1 | 0/1 | 0/1 | 0/1 | 0/1 | 0/1 |
| 0 | 0 | 0 | 0/1 | 0 | 0 | 0/1 |
| 0 | 0/1 | 0/1 | 0/1 | 0/1 | 0 | 0/1 |
| 0 | 0 | 0 | 0 | 0 | 0 | 0/1 |
| 0 | 0 | 0 | 0 | 0 | 0 | 0/1 |
| 0 | 0 | 1/1 | 0 | 1/1 | 0/1 | 0/1 |
| 0 | 0 | 0/1 | 0 | 0 | 0 | 0/1 |
| 0/1 | 0/1 | 0/1 | 0 | 0 | 0/1 | 0/1 |
| 0 | 0 | 0 | 0 | 0 | 0 | 0/1 |
| 0 | 1/1 | 0/1 | 1/1 | 0 | 0/1 | 0/1 |
| 0 | 0 | 0 | 0 | 0 | 0 | 0/1 |

| | | | | | | |
|---|---|---|---|---|---|---|
| 0 | 1/1 | 0 | 0 | 0 | 0/1 | 0/1 |
| 0 | 0 | 0 | 0 | 0 | 0 | 0/1 |
| 0 | 0 | 0 | 0 | 0 | 0 | 0/1 |
| 0 | 0 | 0 | 0 | 0 | 0 | 0/1 |
| 0/1 | 1/1 | 0 | 0/1 | 0 | 0 | 0/1 |
| 0/1 | 1/1 | 0 | 0/1 | 0 | 0 | 0/1 |
| 0/1 | 1/1 | 0 | 0/1 | 0 | 0 | 0/1 |
| 0 | 0 | 0 | 0/1 | 0 | 0 | 0/1 |
| 0/1 | 1/1 | 1/1 | 1/1 | 0 | 0/1 | 0/1 |
| 0 | 0/1 | 0 | 0 | 0 | 0 | 0/1 |
| 0 | 0 | 0 | 0 | 0 | 0 | 0/1 |
| 0 | 0 | 0 | 0 | 0 | 0 | 0/1 |
| 0 | 0 | 0 | 0 | 0 | 0 | 0/1 |
| 0 | 0 | 0 | 0 | 0 | 0/1 | 0/1 |
| 0/1 | 0/1 | 0/1 | 1/1 | 0 | 0/1 | 0/1 |
| 0 | 0 | 0 | 0 | 0 | 0 | 0/1 |
| 0 | 0 | 0 | 0 | 0 | 0 | 0/1 |
| 0 | 1/1 | 1/1 | 0/1 | 0 | 1/1 | 0/1 |
| 0 | 1/1 | 0 | 0 | 0 | 0/1 | 0/1 |
| 0 | 0 | 0 | 0 | 0 | 0 | 0/1 |
| 0 | 0 | 0 | 0 | 0 | 0/1 | 0/1 |
| 1/1 | 0/1 | 0 | 0/1 | 0 | 0 | 0/1 |
| 0 | 0/1 | 0 | 0/1 | 0 | 0 | 0/1 |
| 1/1 | 0/1 | 0 | 0/1 | 0 | 1/1 | 0/1 |
| 0 | 0/1 | 0 | 0 | 0/1 | 0 | 0/1 |
| 1/1 | 0/1 | 0 | 1/1 | 0 | 1/1 | 0/1 |
| 0/1 | 0/1 | 0 | 0/1 | 0 | 0/1 | 0/1 |
| 0/1 | 0/1 | 0 | 0/1 | 0 | 0/1 | 0/1 |
| 1/1 | 0 | 0 | 0/1 | 0 | 1/1 | 0/1 |
| 1/1 | 0/1 | 0 | 0/1 | 0 | 1/1 | 0/1 |
| 1/1 | 0/1 | 0 | 0/1 | 0 | 1/1 | 0/1 |
| 0 | 0/1 | 0 | 0/1 | 0 | 1/1 | 0/1 |
| 0 | 0 | 0 | 0 | 0 | 0 | 0/1 |
| 0 | 0 | 0 | 0 | 0 | 0 | 0/1 |
| 0/1 | 0/1 | 0/1 | 0/1 | 0 | 0/1 | 0/1 |
| 0 | 0 | 0 | 0 | 0 | 0 | 0/1 |

Supplementary Table 2

| B2 | C3 | D4 | E5 | F6 | #Uploaded_variatio | Allele |
|---|---|---|---|---|---|---|
| 0/1 | 1/1 | 1/1 | 1/1 | 1/1 | 1-152883680-A-G | G |
| 0/1 | 1/1 | 1/1 | 1/1 | 1/1 | 1-152883711-G-C | C |
| 0/1 | 1/1 | 1/1 | 1/1 | 1/1 | 1-152884111-G-A | A |
| 0/1 | 1/1 | 1/1 | 1/1 | 1/1 | 1-153012765-T-C | C |
| 0/1 | 1/1 | 1/1 | 1/1 | 1/1 | 1-153363105-T-C | C |
| 0/1 | 1/1 | 1/1 | 1/1 | 1/1 | 1-153588340-C-T | T |
| 0/1 | 1/1 | 1/1 | 1/1 | 1/1 | 1-154401972-T-C | C |
| 0/1 | 1/1 | 1/1 | 1/1 | 1/1 | 1-154426970-A-C | C |
| 0/1 | 1/1 | 1/1 | 1/1 | 1/1 | 1-154528053-G-A | A |
| 0/1 | 1/1 | 1/1 | 1/1 | 1/1 | 1-154544651-G-T | T |
| 0/1 | 1/1 | 1/1 | 1/1 | 1/1 | 1-154580458-C-A | A |
| 0/1 | 1/1 | 1/1 | 1/1 | 1/1 | 1-154899040-G-A | A |
| 0/1 | 1/1 | 1/1 | 1/1 | 1/1 | 1-155033317-T-C | C |
| 0/1 | 1/1 | 1/1 | 1/1 | 1/1 | 1-155034632-C-G | G |
| 0/1 | 1/1 | 1/1 | 1/1 | 1/1 | 1-155105882-T-G | G |
| 0/1 | 1/1 | 1/1 | 1/1 | 1/1 | 1-155106550-A-G | G |
| 0/1 | 1/1 | 1/1 | 1/1 | 1/1 | 1-155108287-G-A | A |
| 0/1 | 1/1 | 1/1 | 1/1 | 1/1 | 1-155142210-A-G | G |
| 0/1 | 1/1 | 1/1 | 1/1 | 1/1 | 1-155142229-G-A | A |
| 0/1 | 1/1 | 1/1 | 1/1 | 1/1 | 1-155148781-G-A | A |
| 0/1 | 1/1 | 1/1 | 1/1 | 1/1 | 1-155149718-G-A | A |
| 0/1 | 1/1 | 1/1 | 1/1 | 1/1 | 1-155162067-C-T | T |
| 0/1 | 1/1 | 1/1 | 1/1 | 1/1 | 1-156025096-C-T | T |
| 0/1 | 1/1 | 1/1 | 1/1 | 1/1 | 1-156202173-A-G | G |
| 0/1 | 1/1 | 1/1 | 1/1 | 1/1 | 1-156206158-C-T | T |
| 0/1 | 1/1 | 1/1 | 1/1 | 1/1 | 1-156213921-C-T | T |
| 0/1 | 1/1 | 1/1 | 1/1 | 1/1 | 1-156230206-C-A | A |
| 0/1 | 1/1 | 1/1 | 1/1 | 1/1 | 1-156261491-G-C | C |
| 0/1 | 1/1 | 1/1 | 1/1 | 1/1 | 1-156498673-T-A | A |
| 0/1 | 1/1 | 1/1 | 1/1 | 1/1 | 1-156499969-T-C | C |
| 0/1 | 1/1 | 1/1 | 1/1 | 1/1 | 1-156502870-C-T | T |
| 0/1 | 1/1 | 1/1 | 1/1 | 1/1 | 1-156641537-C-T | T |
| 0/1 | 1/1 | 1/1 | 1/1 | 1/1 | 1-156713558-G-A | A |
| 0/1 | 1/1 | 1/1 | 1/1 | 1/1 | 1-156761618-C-T | T |
| 0/1 | 1/1 | 1/1 | 1/1 | 1/1 | 1-156834364-G-A | A |
| 0/1 | 1/1 | 1/1 | 1/1 | 1/1 | 1-156882950-C-T | T |
| 0/1 | 1/1 | 1/1 | 1/1 | 1/1 | 1-156882996-G-C | C |
| 0/1 | 1/1 | 1/1 | 1/1 | 1/1 | 1-156883617-G-A | A |
| 0/1 | 1/1 | 1/1 | 1/1 | 1/1 | 1-157550066-T-C | C |
| 0/1 | 1/1 | 1/1 | 1/1 | 1/1 | 1-157768030-G-C | C |
| 0/1 | 1/1 | 1/1 | 1/1 | 1/1 | 1-157771880-C-T | T |
| 0/1 | 1/1 | 1/1 | 1/1 | 1/1 | 1-157802877-T-C | C |

| | | | | | | |
|---|---|---|---|---|---|---|
| 0/1 | 1/1 | 1/1 | 1/1 | 1/1 | 1-157802979-C-T | T |
| 0/1 | 1/1 | 1/1 | 1/1 | 1/1 | 1-157806006-G-A | A |
| 0/1 | 1/1 | 1/1 | 1/1 | 1/1 | 1-157894802-C-T | T |
| 0/1 | 1/1 | 1/1 | 1/1 | 1/1 | 1-158325891-T-C | C |
| 0/1 | 1/1 | 1/1 | 1/1 | 1/1 | 1-158368964-C-T | T |
| 0/1 | 1/1 | 1/1 | 1/1 | 1/1 | 1-158368974-A-G | G |
| 0/1 | 1/1 | 1/1 | 1/1 | 1/1 | 1-158369064-A-G | G |
| 0/1 | 1/1 | 1/1 | 1/1 | 1/1 | 1-158577167-A-T | T |
| 0/1 | 1/1 | 1/1 | 1/1 | 1/1 | 1-159505297-C-T | T |
| 0/1 | 1/1 | 1/1 | 1/1 | 1/1 | 1-159850401-C-G | G |
| 0/1 | 1/1 | 1/1 | 1/1 | 1/1 | 1-159897327-C-T | T |
| 0/1 | 1/1 | 1/1 | 1/1 | 1/1 | 1-159897754-C-T | T |
| 0/1 | 1/1 | 1/1 | 1/1 | 1/1 | 1-160011455-T-C | C |
| 0/1 | 1/1 | 1/1 | 1/1 | 1/1 | 1-160106531-G-A | A |
| 0/1 | 1/1 | 1/1 | 1/1 | 1/1 | 1-160134205-C-T | T |
| 0/1 | 1/1 | 1/1 | 1/1 | 1/1 | 1-160137300-A-C | C |
| 0/1 | 1/1 | 1/1 | 1/1 | 1/1 | 1-160146173-T-C | C |
| 0/1 | 1/1 | 1/1 | 1/1 | 1/1 | 1-160147502-T-G | G |
| 0/1 | 1/1 | 1/1 | 1/1 | 1/1 | 1-160147510-G-A | A |
| 0/1 | 1/1 | 1/1 | 1/1 | 1/1 | 1-161049499-G-A | A |
| 0/1 | 1/1 | 1/1 | 1/1 | 1/1 | 1-161088292-A-G | G |
| 0/1 | 1/1 | 1/1 | 1/1 | 1/1 | 1-161088583-T-G | G |
| 0/1 | 1/1 | 1/1 | 1/1 | 1/1 | 1-161092761-A-G | G |
| 0/1 | 1/1 | 1/1 | 1/1 | 1/1 | 1-161094190-A-G | G |
| 0/1 | 1/1 | 1/1 | 1/1 | 1/1 | 1-161126975-C-G | G |
| 0/1 | 1/1 | 1/1 | 1/1 | 1/1 | 1-161136843-C-G | G |
| 0/1 | 1/1 | 1/1 | 1/1 | 1/1 | 1-161139738-G-A | A |
| 0/1 | 1/1 | 1/1 | 1/1 | 1/1 | 1-161143801-G-A | A |
| 0/1 | 1/1 | 1/1 | 1/1 | 1/1 | 1-161145005-C-T | T |
| 0/1 | 1/1 | 1/1 | 1/1 | 1/1 | 1-161161284-G-C | C |
| 0/1 | 1/1 | 1/1 | 1/1 | 1/1 | 1-161172233-C-A | A |
| 0/1 | 1/1 | 1/1 | 1/1 | 1/1 | 1-161179877-G-A | A |
| 0/1 | 1/1 | 1/1 | 1/1 | 1/1 | 1-161495040-C-T | T |
| 0/1 | 1/1 | 1/1 | 1/1 | 1/1 | 1-161495117-C-T | T |
| 0/1 | 1/1 | 1/1 | 1/1 | 1/1 | 1-161495477-A-T | T |
| 0/1 | 1/1 | 1/1 | 1/1 | 1/1 | 1-161496134-C-G | G |

Supplementary Table 2

| Consequence | SYMBOL | Protein_position | Amino_acids | Codons |
|---|---|---|---|---|
| synonymous_variant | IVL | 469 | Q | caA/caG |
| missense_variant | IVL | 480 | V/L | Gtg/Ctg |
| 3_prime_UTR_variant | IVL | - | - | - |
| missense_variant | SPRR2D | 20 | T/A | Acg/Gcg |
| intron_variant | S100A8 | - | - | - |
| 5_prime_UTR_variant | S100A14 | - | - | - |
| intron_variant | IL6R | - | - | - |
| missense_variant | IL6R | 358 | D/A | gAt/gCt |
| intron_variant | UBE2Q1 | - | - | - |
| intron_variant | CHRNB2 | - | - | - |
| intron_variant | ADAR | - | - | - |
| intron_variant | PMVK | - | - | - |
| intron_variant | ADAM15 | - | - | - |
| intron_variant | ADAM15 | - | - | - |
| intron_variant | EFNA1 | - | - | - |
| 3_prime_UTR_variant | EFNA1 | - | - | - |
| intron_variant | SLC50A1 | - | - | - |
| intron_variant | KRTCAP2 | - | - | - |
| intron_variant | KRTCAP2 | - | - | - |
| intron_variant | TRIM46 | - | - | - |
| synonymous_variant | TRIM46 | 287 | T | acG/acA |
| synonymous_variant | MUC1 | 22 | T | acG/acA |
| synonymous_variant | LAMTOR2 | 37 | Y | taC/taT |
| missense_variant | PMF1-BGLAP | 75 | Q/R | cAa/cGa |
| synonymous_variant | PMF1-BGLAP | 129 | G | ggC/ggT |
| missense_variant | PAQR6 | 263 | E/K | Gag/Aag |
| intron_variant | SMG5 | - | - | - |
| 3_prime_UTR_variant | TMEM79 | - | - | - |
| intron_variant | IQGAP3 | - | - | - |
| synonymous_variant | IQGAP3 | 1444 | L | ctA/ctG |
| synonymous_variant | IQGAP3 | 1335 | T | acG/acA |
| missense_variant | NES | 815 | V/I | Gta/Ata |
| missense_variant | HDGF | 217 | P/L | cCc/cTc |
| intron_variant | PRCC | - | - | - |
| intron_variant | NTRK1 | - | - | - |
| intron_variant | PEAR1 | - | - | - |
| synonymous_variant | PEAR1 | 811 | P | ccG/ccC |
| intron_variant | PEAR1 | - | - | - |
| intron_variant | FCRL4 | - | - | - |
| missense_variant | FCRL1 | 345 | S/R | agC/agG |
| synonymous_variant | FCRL1 | 237 | P | ccG/ccA |
| intron_variant | CD5L | - | - | - |

| | | | | |
|---|---|---|---|---|
| splice_region_variant,intro | CD5L | - | - | - |
| intron_variant | CD5L | - | - | - |
| intergenic_variant | - | - | - | - |
| synonymous_variant | CD1E | 300 | H | caT/caC |
| missense_variant | OR10T2 | 98 | C/Y | tGt/tAt |
| missense_variant | OR10T2 | 95 | F/L | Ttc/Ctc |
| missense_variant | OR10T2 | 65 | F/L | Ttc/Ctc |
| synonymous_variant | OR10Z1 | 313 | G | ggA/ggT |
| synonymous_variant | OR10J5 | 167 | P | ccG/ccA |
| synonymous_variant | CCDC19 | 329 | L | ctG/ctC |
| intron_variant | IGSF9 | - | - | - |
| intron_variant | IGSF9 | - | - | - |
| missense_variant | KCNJ10 | 290 | T/A | Acc/Gcc |
| intron_variant | ATP1A2 | - | - | - |
| synonymous_variant | ATP1A4 | 346 | A | gcC/gcT |
| intron_variant | ATP1A4 | - | - | - |
| intron_variant | ATP1A4 | - | - | - |
| intron_variant | ATP1A4 | - | - | - |
| intron_variant | ATP1A4 | - | - | - |
| missense_variant | PVRL4 | 107 | P/L | cCc/cTc |
| intron_variant | NIT1 | - | - | - |
| missense_variant | NIT1 | 4 | F/V | Ttc/Gtc |
| intron_variant | DEDD | - | - | - |
| synonymous_variant | DEDD | 21 | H | caT/caC |
| intron_variant | UFC1 | - | - | - |
| intron_variant | PPOX | - | - | - |
| missense_variant | PPOX | 304 | R/H | cGt/cAt |
| synonymous_variant | B4GALT3 | 176 | N | aaC/aaT |
| synonymous_variant | B4GALT3 | 89 | S | tcG/tcA |
| missense_variant | ADAMTS4 | 720 | P/A | Cct/Gct |
| missense_variant | NDUFS2 | 20 | P/T | Cct/Act |
| intron_variant | NDUFS2 | - | - | - |
| missense_variant | HSPA6 | 198 | L/F | Ctc/Ttc |
| synonymous_variant | HSPA6 | 223 | A | gcC/gcT |
| synonymous_variant | HSPA6 | 343 | T | acA/acT |
| missense_variant | HSPA6 | 562 | D/E | gaC/gaG |

Supplementary Table 2

| Existing_variation | STRAND | SIFT | SIFT_score | PolyPhen | PolyPhen_score |
|---|---|---|---|---|---|
| rs7535306,COSM34 | 1 | - | | - | |
| rs7545520 | 1 | tolerated_low_( | 0.39 | benign | 0 |
| rs913996 | 1 | - | | - | |
| rs1846857 | -1 | tolerated_low_( | 0.23 | benign | 0.003 |
| rs3795391 | -1 | - | | - | |
| rs11548103,CR0921 | -1 | - | | - | |
| rs6694817 | 1 | - | | - | |
| rs2228145,CM03473 | 1 | tolerated | 0.08 | benign | 0.02 |
| rs56047170 | -1 | - | | - | |
| rs4845378 | 1 | - | | - | |
| rs58655370 | -1 | - | | - | |
| - | -1 | - | | - | |
| rs11264304 | 1 | - | | - | |
| rs45444697 | 1 | - | | - | |
| rs4971066 | 1 | - | | - | |
| rs9297 | 1 | - | | - | |
| rs11264333 | 1 | - | | - | |
| rs4421576 | -1 | - | | - | |
| rs4276914 | -1 | - | | - | |
| rs4971059 | 1 | - | | - | |
| rs3814316 | 1 | - | | - | |
| rs4072037 | -1 | - | | - | |
| rs7541 | 1 | - | | - | |
| rs1052053 | 1 | tolerated_low_( | 1 | benign | 0 |
| rs62001899 | 1 | - | | - | |
| rs7513351 | -1 | deleterious_low | 0 | benign | 0.015 |
| rs2287024 | -1 | - | | - | |
| rs3795728 | 1 | - | | - | |
| rs1171564 | -1 | - | | - | |
| rs1171566 | -1 | - | | - | |
| rs4661044 | -1 | - | | - | |
| rs951781 | -1 | tolerated | 0.6 | benign | 0 |
| rs4399146 | -1 | deleterious_low | 0 | benign | 0 |
| rs2644615 | 1 | - | | - | |
| rs72698666 | 1 | - | | - | |
| rs61813832 | 1 | - | | - | |
| rs822441 | 1 | - | | - | |
| rs61813833 | 1 | - | | - | |
| rs10489673 | -1 | - | | - | |
| rs149687405 | -1 | tolerated | 0.06 | benign | 0.36 |
| rs4971154 | -1 | - | | - | |
| rs2765500 | -1 | - | | - | |

| | | | | | |
|---|---|---|---|---|---|
| rs2261295 | -1 | - | | - | |
| rs2765503 | -1 | - | | - | |
| rs114568699 | - | - | | - | |
| rs61734679 | 1 | - | | - | |
| rs61818749,COSM1 | -1 | deleterious | 0 | probably_dam | 0.999 |
| rs61818750 | -1 | tolerated | 0.53 | benign | 0.05 |
| rs41488350 | -1 | tolerated | 0.22 | benign | 0.148 |
| rs2427808 | 1 | - | | - | |
| rs4656837 | -1 | - | | - | |
| rs7550537 | -1 | - | | - | |
| rs2153208 | -1 | - | | - | |
| rs80070763 | -1 | - | | - | |
| - | -1 | deleterious | 0 | probably_dam | 0.999 |
| rs12083034 | 1 | - | | - | |
| rs11265338 | 1 | - | | - | |
| rs4656889 | 1 | - | | - | |
| rs881291 | 1 | - | | - | |
| rs731852 | 1 | - | | - | |
| rs731853 | 1 | - | | - | |
| rs78105657 | -1 | tolerated | 0.34 | probably_dam | 0.999 |
| rs138523655,COSM | 1 | - | | - | |
| rs41270017 | 1 | tolerated_low_ | 0.27 | benign | 0 |
| rs17389237 | -1 | - | | - | |
| rs1135783 | -1 | - | | - | |
| rs12031437 | 1 | - | | - | |
| rs2301287 | 1 | - | | - | |
| rs36013429 | 1 | tolerated | 0.36 | benign | 0 |
| rs3813619 | -1 | - | | - | |
| rs3813620 | -1 | - | | - | |
| rs41270041 | -1 | tolerated | 0.23 | benign | 0.01 |
| rs11538340 | 1 | tolerated_low_ | 0.29 | benign | 0.003 |
| rs4656994 | 1 | - | | - | |
| rs1079109 | 1 | deleterious_lov | 0 | probably_dam | 1 |
| rs41297716,COSM3 | 1 | - | | - | |
| - | 1 | - | | - | |
| rs753856 | 1 | tolerated_low_ | 0.32 | possibly_dam | 0.469 |
