## Supplemental table 3 for "Identification and functional characterization of two novel mutations in *KCNJ10* and *PI4KB* in SeSAME syndrome without electrolyte imbalance"

**Supplementary Table 3: Variants (n=7)  shortlisted based on assessing the allele freq**

| Chr | Start | Ref | Alt | Func.refGene | Gene.refGer | ExonicFunc.refGene |
|---|---|---|---|---|---|---|
| 1 | 151288779 | T | C | exonic | PI4KB | nonsynonymous SNV |
| 1 | 151787451 | C | T | exonic | RORC | nonsynonymous SNV |
| 1 | 152324002 | T | C | exonic | FLG2 | nonsynonymous SNV |
| 1 | 157768030 | G | C | exonic | FCRL1 | nonsynonymous SNV |
| 1 | 160011455 | T | C | exonic | KCNJ10 | nonsynonymous SNV |
| 1 | 161049499 | G | A | exonic | PVRL4 | nonsynonymous SNV |
| 1 | 161088292 | A | G | exonic | NIT1 | nonsynonymous SNV |

**quencies (MAF<0.01) in 1KG_all and ExAC_all. The below variants are the only those wh**

| 1000g2015aug_all | ExAC_ALL | SIFT_score | SIFT_pred | Polyphen2_HDIV_score |
|---|---|---|---|---|
| 0 | 0 | 0.039 | D | 0.215 |
| 0.00239617 | 0.0035 | 0.322 | T | 0.001 |
| 0.00479233 | 0.0068 | 0.064 | T | 0.015 |
| 0.00359425 | 0.0036 | 0.097 | T | 0.893 |
| 0 | 0 | 0 | D | 1 |
| 0.00419329 | 0.0052 | 0.556 | T | 1 |
| 0.000998403 | 0.0051 | 0.035 | D | 0 |

**ıich were HET in all unaffected parents and HOM in all affected siblings.**

| Polyphen2_HDIV_pred |
|---|
| B |
| B |
| B |
| P |
| D |
| D |
| B |
